# Macropinocytosis enables metabolic recycling of extracellular DNA in cancer cells

**DOI:** 10.1101/2025.10.22.684010

**Authors:** Wen-Hsuan Yang, Olivia T. Stamatatos, Evdokia Michalopoulou, Ava Sann, Paolo Cifani, Linda Van Aelst, Justin R. Cross, Tse-Luen Wee, Michael J. Lukey

## Abstract

Avid nutrient consumption is a metabolic hallmark of cancer and leads to regional depletion of key metabolites within the tumor microenvironment (TME). Cancer cells consequently employ diverse strategies to acquire the fuels needed for growth, including bulk uptake of the extracellular medium by macropinocytosis. Here, we show that breast and pancreatic cancer cells macropinocytically internalize extracellular DNA (exDNA), an abundant component of the TME, and deliver it to lysosomes for degradation. This provides a supply of nucleotides that sustains growth when *de novo* biosynthesis is impaired by glutamine restriction or pharmacological blockade. Mechanistically, this process is dependent on the non-redundant lysosomal equilibrative nucleoside transporter SLC29A3 (ENT3), which mediates the export of nucleosides from the lysosomal lumen into the cytosol. Accordingly, genetic ablation of SLC29A3 or pharmacological disruption of lysosomal function prevents exDNA scavenging and potently sensitizes breast tumors to antimetabolite chemotherapy *in vivo*. These findings reveal a previously unrecognized nutrient acquisition pathway through which cancer cells recycle exDNA into metabolic building blocks and highlight SLC29A3 as a mediator of metabolic flexibility and a potential target to improve chemotherapy response.

## Introduction

Avid nutrient consumption is a conserved hallmark of cancer metabolism and is needed to fuel the biosynthetic and bioenergetic pathways that sustain tumor growth^1,2^. However, intense cellular competition for available resources within the tumor microenvironment (TME), compounded by the disorganized, occluded, and leaky nature of the tumor vasculature, results in regions of severe intratumoral nutrient depletion^3–6^. Cancer cells consequently employ a variety of strategies to acquire essential metabolites, including transporter-mediated uptake, receptor-mediated endocytosis, entosis, and macropinocytosis, with the mode of acquisition being influenced by the metabolic state, cell lineage, and oncogenotype^7^.

Macropinocytosis is an evolutionarily conserved form of clathrin-independent endocytosis that involves the non-selective uptake of extracellular fluid in vesicles termed macropinosomes^8,9^. Formation of macropinosomes occurs at phosphatidylinositol-3-phosphate (PIP3) domains on the inner leaflet of the plasma membrane and is driven by a ring of actin polymerization that leads to encapsulation of the bulk extracellular medium^10^. Following a macropinosome’s scission, Rab GTPases coordinate its trafficking through the endolysosomal system, culminating in macropinosome-lysosome fusion and cargo degradation^11,12^. Although macropinocytosis is normally tightly regulated by growth factor signaling, specific oncogenic lesions including activating *KRAS* and *PIK3CA* mutations can drive constitutive macropinocytic activity in cancer cells^13–15^. It has recently emerged that the macropinocytic uptake and lysosomal digestion of extracellular protein is an important amino acid supply mechanism in nutrient-deprived tumors including *KRAS*-mutant pancreatic ductal adenocarcinoma (PDAC) and *PIK3CA*-mutant breast cancer^12–14,16,17^.

One of the major anabolic pathways drawing on supplies of biosynthetic precursors in tumors is *de novo* nucleotide biosynthesis, which is highly upregulated during malignant transformation and has been described as a pan-cancer metabolic dependency^18^. Nucleotide biosynthesis requires a ribose sugar moiety, typically derived from glucose, as well as one-carbon units, the amino acids glycine, aspartate, and glutamine, and oxidizing power and energy in the form of nicotinamide adenine dinucleotide (NAD^+^) and ATP, respectively. The amide group of glutamine is the obligate nitrogen source for 5 separate reactions in purine and pyrimidine biosynthesis, making this amino acid a strict metabolic requirement for *de novo* nucleotide biosynthesis^18,19^. While non-proliferative cells typically maintain their nucleotide pools via purine and pyrimidine salvage pathways, nucleotide levels increase ∼5-10-fold during proliferation and are elevated even further in cancer cells, necessitating *de novo* synthesis^18,20,21^. Aligned with the key importance of nucleotide biosynthesis in cancer, antimetabolite inhibitors of this pathway such as methotrexate, 5-flurouracil, and mercaptopurine (6-MP) are among the most widely used drugs in clinical oncology. However, their application is restricted by dose-limiting toxicity and the eventual acquisition of drug resistance^22^.

Here, we report that breast cancer and PDAC cells can overcome their dependence on glutamine-derived nitrogen and *de novo* nucleotide biosynthesis by employing macropinocytosis to scavenge extracellular DNA (exDNA). exDNA is ubiquitously present in mammalian tissues and blood plasma and its abundance increases in a spectrum of disease states including cancer, with sources including apoptotic and necrotic cells, neutrophils undergoing NETosis, and secreted mitochondrial DNA^23–25^. We show that exDNA-laden macropinosomes are trafficked to lysosomes, where the equilibrative nucleoside transporter SLC29A3 (ENT3) is required to process the cargo. Although SLC29A3 is dispensable for cancer cell proliferation in nutrient-rich environments, it becomes essential when cancer cells rely on exDNA consumption, as occurs during glutamine starvation or treatment with chemotherapies targeting nucleotide biosynthesis. As such, we find that genetic depletion of SLC29A3 potently sensitizes breast tumors to antipurine treatment, effectively eliminating tumor growth. Our findings thus reveal a new pathway for nitrogen scavenging in nutrient-deprived cancer cells and identify potential therapeutic approaches to maximize the efficacy of antimetabolite chemotherapy.

## Results

### Cancer cells consume exDNA by macropinocytosis and traffic it to lysosomes

Circulating exDNA is widely elevated in patients with cancer and is used as a diagnostic and prognostic tool in clinical oncology^23,25^. The microenvironment of many solid tumors is also highly enriched with exDNA, which exists almost exclusively in an exposed non-membrane-protected form and is derived primarily from dying cells^23,25–28^. To probe for exDNA in the TME of breast cancer and PDAC, we performed immunofluorescence analysis of non-permeabilized fresh tumor tissue sections from human patients and mouse models using a monoclonal antibody that specifically binds DNA. This yielded an extracellular signal (Fig. 1a,b and Extended Data Fig. 1a,b), with a gradient that increased from the tumor periphery to the necrotic core (Fig. 1c and Extended Data Fig. 1c). Supporting on-target binding of the antibody, treatment of tumor slices with deoxyribonuclease I (DNase I) sharply depleted the exDNA signal (Fig. 1d), and additional control experiments confirmed that the antibody recognizes gel-purified DNA and that this signal is also lost upon DNase I treatment (Extended Data Fig. 1d). We observed that neutrophil extracellular traps (NETs) are present throughout the tumor samples, identified by the marker citrullinated histone H3 (Fig. 1e), but these do not account for all of the exDNA signal.

**Fig. 1:**
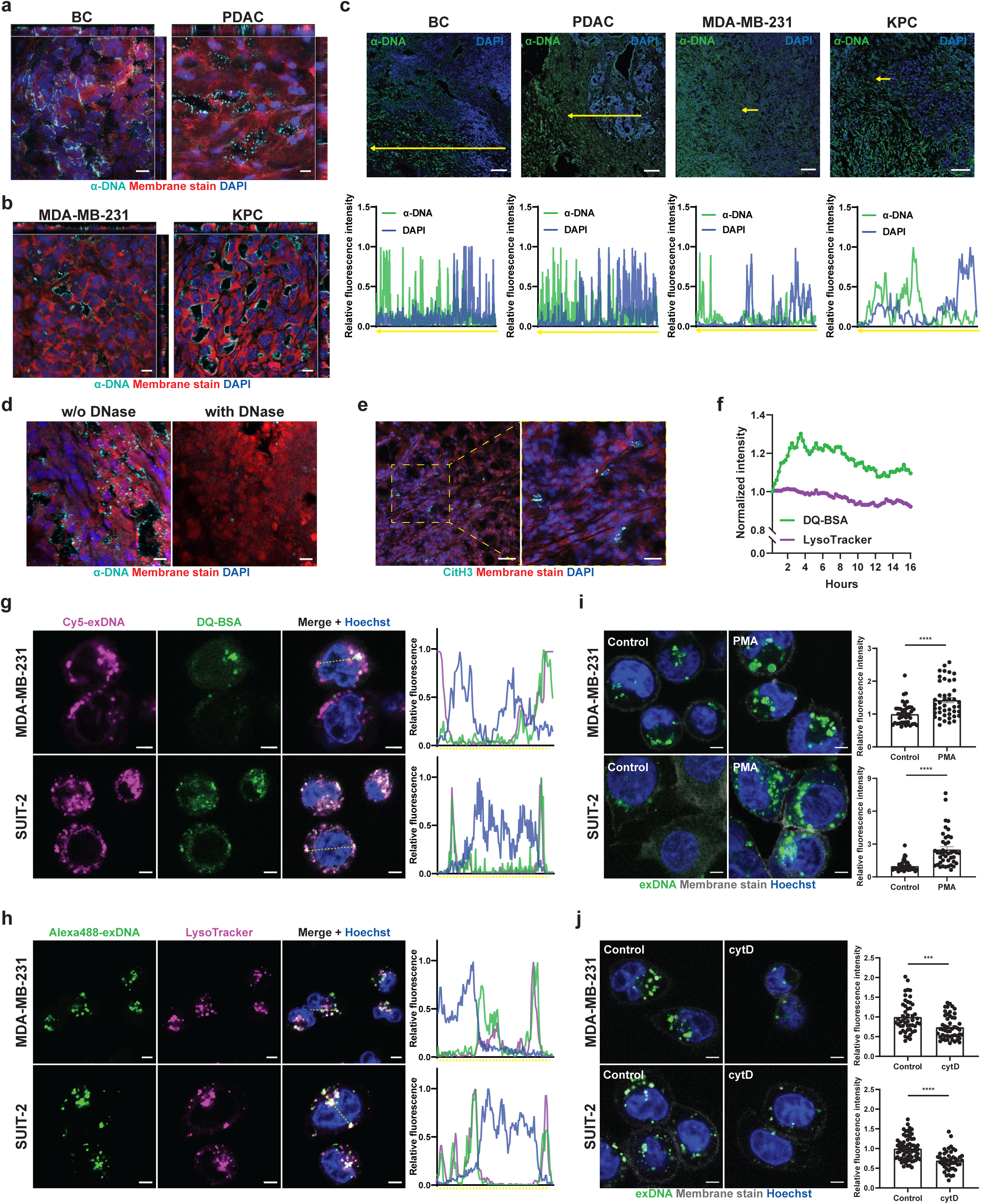
Cancer cells consume exDNA by macropinocytosis and traffic it to lysosomes. **a**, Z-stack confocal images of human tumors. Left: Breast cancer (BC) tumor; Right: Pancreatic ductal adenocarcinoma (PDAC) tumor. The samples were stained with an anti-DNA antibody (turquoise), membrane stain (red) for tissue boundaries, and DAPI (blue) for nuclei. Scale bar: 10 µm. **b**, Z-stack confocal images of human MDA-MB-231 breast cancer xenografts (left) and spontaneous PDAC tumors from KPC mice (right) with an anti-DNA antibody (turquoise), membrane stain (red), and DAPI (blue). Scale bar, 10 µm. **c**, Top: Representative confocal fluorescence microscopy images of human BC, human PDAC, orthotopic breast cancer xenografts (MDA-MB-231), and spontaneous PDAC tumors from KPC mice. DNA was visualized using an anti-DNA-specific antibody (green) and nuclei were counterstained with DAPI (blue). The arrow indicates the direction from the tumor periphery toward the core. Images acquired using a 20× objective and 5×5 tile scanning. Scale bars: 100 µm (BC and KPC) and 200 µm (MDA-MB-231 and PDAC). Bottom: Confocal immunofluorescence analysis with relative pixel intensity. **d**, Confocal fluorescence microscopy images of MDA-MB-231 tumor slides treated with or without DNase I, showing anti-DNA antibody staining (turquoise), membrane staining (red), and DAPI nuclear staining (blue). Scale bar, 10 µm. **e**, Confocal fluorescence microscopy of MDA-MB-231 xenograft tumor sections showing citrullinated histone H3 (turquoise), membrane staining (red), and DAPI nuclear staining (blue). Scale bars: 50 µm (left) and 20 µm (right, zoom-in). **f**, Macropinocytosis dynamics in MDA-MB-231 cells monitored using Lysotracker Red and DQ-BSA, with images acquired every 15 minutes on a ZEISS Lattice Lightsheet 7. Fluorescence intensities were normalized to baseline, n = 5. **g**, Confocal immunofluorescence analysis of Cy5-exDNA (magenta) and DQ-BSA (green) in MDA-MB-231 breast cancer (upper row) and SUIT-2 PDAC (lower row) cell lines. Nuclei were stained with Hoechst 33342 (blue). Images are shown individually and merged, with corresponding fluorescence quantification plots. Scale bars: 5 µm. **h**, Confocal immunofluorescence analysis of Alexa488-exDNA (green) and LysoTracker Red (magenta) in MDA-MB-231 and SUIT-2 cells. Nuclei were stained with Hoechst 33342 (blue). Images are shown individually and merged, with corresponding fluorescence quantification plots. Scale bars: 5 µm. **i**, Left, confocal immunofluorescence images of MDA-MB-231 and SUIT-2 cell lines treated with or without 250 nM 12-myristate 13-acetate (PMA) and 2.5 µM fluorophore-labeled exDNA overnight, showing exDNA uptake (green). Cell boundaries were stained with membrane dye (gray), and nuclei with Hoechst 33342 (blue). Scale bars: 5 μm. Fluorescence intensity was quantified by measuring vesicle mean intensity using the Cell tool in the Imaris analysis platform. Statistical analysis was performed using two-tailed unpaired Student’s *t* tests. **j**, Left, confocal immunofluorescence images of MDA-MB-231 and SUIT-2 cells treated with or without 25 nM cytD and Alexa488-labeled exDNA overnight to assess exDNA uptake (green). Cell boundaries were stained with a membrane dye (gray), and nuclei were counterstained with Hoechst 33342 (blue). Scale bars: 5 μm. Fluorescence intensity was quantified by measuring vesicle mean intensity using the Imaris Cell tool. Statistical significance was determined using two-tailed unpaired Student’s *t* tests.

Breast cancer and PDAC cells with oncogenic *PIK3CA* or *KRAS* mutations typically have high constitutive macropinocytic activity, permitting the use of extracellular protein as a nutrient source in culture and *in vivo*^16,17,29^. Consistent with previous findings^16,30^, live-cell imagining revealed that MDA-MB-231 breast cancer cells avidly consume the labeled protein DQ-BSA and traffic it to lysosomes, where it becomes fluorescent upon lysosomal digestion (Fig. 1f and Supplementary Video 1)^31,32^. We hypothesized that exDNA might also be consumed by macropinocytosis in these cells. To test this, we supplied 30-mer random oligonucleotides labeled with a 5’ cyanine 5 (Cy5) fluorophore and with a GC content of 41%, corresponding to the average GC content of the human genome^33^, to cultured breast cancer and PDAC cells, along with DQ-BSA for 16 h. Confocal fluorescence microscopy revealed robust cellular uptake of Cy5-DNA, and colocalization analysis showed substantial overlap between the Cy5-DNA signal and the lysosomal DQ-BSA signal (Fig. 1g and Extended Data Fig. 1e). To obtain further evidence that exDNA can be consumed and trafficked to lysosomes, we also supplied cells with Alexa488-DNA and co-stained with the lysosomal dye LysoTracker Red. Again, labeled exDNA was taken up by cancer cells and co-localized with lysosomes (Fig. 1h and Extended Data Fig. 1f).

We next addressed whether cancer cell consumption of exDNA occurs via macropinocytosis, by applying pharmacological activators or inhibitors of this process. Treatment with the diacylglycerol mimetic phorbol 12-myristate 13-acetate (PMA), a stimulator of macropinocytosis^34^, resulted in increased uptake of fluorophore-labeled exDNA in all examined breast cancer and PDAC cell lines (Fig. 1i and Extended Data Fig. 1g). Cytochalasin D (cytD) is a potent inhibitor of actin polymerization that blocks both macropinocytosis and phagocytosis but has additional biological effects when used at higher concentrations^35^. We determined the IC_50_ of cytD against breast cancer and PDAC cells (∼150– 600 nM) and selected a subtoxic concentration of 25 nM, which had no effect on proliferation under nutrient-rich control conditions (Extended Data Fig. 1h). Treatment of cells with this very low dose of cytD suppressed the consumption of exDNA (Fig. 1j and Extended Data Fig. 1i). Similarly, treatment with a subtoxic concentration of the structurally and mechanistically distinct macropinocytosis inhibitor, ethyl-isopropyl amiloride (EIPA), which lowers submembranous pH to prevent activation of the small GTPases that drive macropinosome formation, robustly decreased cellular uptake of exDNA (Extended Data Fig. 1j,k).

### Macropinocytosis of exDNA overcomes cancer cell dependence on glutamine-derived nitrogen

The glutamine requirements of proliferative cells frequently exceed their capacity to synthesize this amino acid *de novo*, resulting in dependence on exogenous sources^36,37^. Major roles of glutamine in intermediary metabolism include the supply of carbon to the tricarboxylic acid (TCA) cycle, and of nitrogen and/or carbon to *de novo* nucleotide, non-essential amino acid (NEAA), glucosamine, and NAD^+^ biosynthesis (Fig. 2a)^38^. Glutamine-mediated TCA cycle anaplerosis is initiated by the deamidation of glutamine to glutamate, a reaction catalyzed by mitochondrial glutaminase (GLS)^39^. In turn, glutamate is converted into the TCA cycle intermediate α-ketoglutarate (α-KG) by glutamate dehydrogenase (GLUD) or a family of transaminases (Extended Data Fig. 2a)^36^. Alternative anaplerotic substrates such as pyruvate and fatty acids, are often readily available *in vivo*, which has confounded efforts to target GLS for cancer therapy^37^. In contrast to this inherent flexibility in carbon supply, the amide group of glutamine is an obligate nitrogen source for *de novo* purine and pyrimidine biosynthesis, making glutamine supply a strict metabolic requirement for these pathways (Fig. 2a,b and Extended Data Fig. 2a,b).

**Fig. 2:**
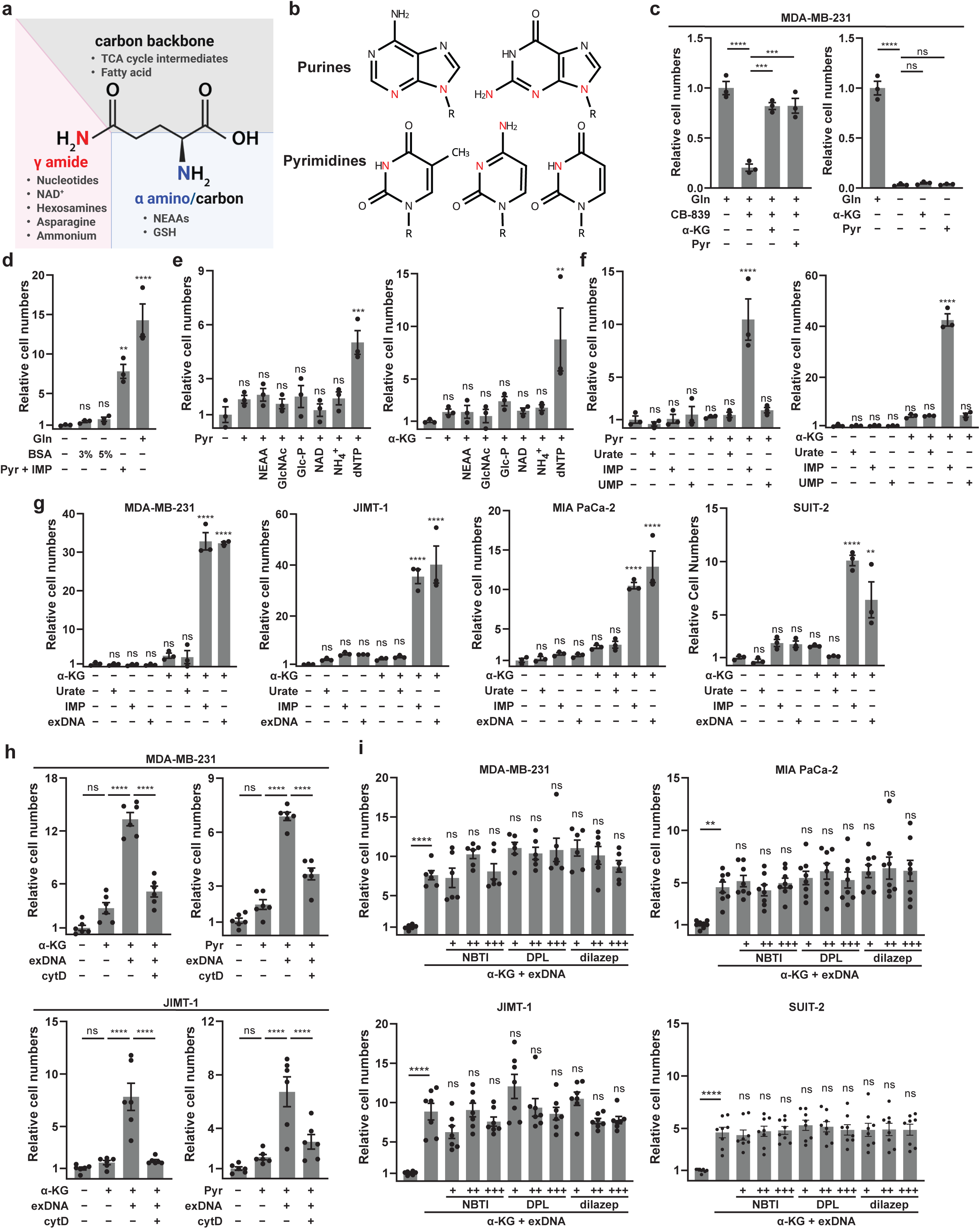
Macropinocytosis of exDNA overcomes cancer cell dependence on glutamine-derived nitrogen. **a**, Overview of cellular metabolic fates of glutamine. **b**, Chemical structures of nucleic acids. The glutamine amide-derived nitrogen is highlighted in red. **c**, MDA-MB-231 cells were treated for 6 days with 0.1 μM CB-839 or glutamine deprivation, with or without 1 mM α-ketoglutarate (α-KG) or pyruvate (Pyr). Cell counts were normalized to the RPMI control and are presented as mean ± SEM, n = 3. **d**, MDA-MB-231 cells for 7 days in glutamine-free medium supplemented with either 3% or 5% BSA, with 1 mM Pyr plus 200 µM inosine monophosphate (IMP) or with 2 mM glutamine (Gln). Cell density was quantified by crystal violet staining and normalized to the glutamine-free control. Statistical significance was assessed by one-way ANOVA against the glutamine-free control. Data are shown as mean ± SEM from three independent replicates. **e**, MDA-MB-231 cells were treated for 10 days under glutamine-free conditions supplemented with 0.2 mM each of the following: non-essential amino acids (NEAA: glycine, alanine, asparagine, aspartate, glutamate, proline, serine), N-acetyl-D-glucosamine (GlcNAc), D-glucosamine 6-phosphate (Glc-P), nicotinamide adenine dinucleotide (NAD^+^), ammonium (NH₄⁺), or mixed dNTPs. Treatments were performed with or without 1 mM Pyr or α-KG. Cell densities were normalized to the glutamine-free control and are presented as mean ± SEM, n = 3. **f**, MDA-MB-231 cells were treated with 200 µM urate, IMP, or UMP ± 1 mM α-KG or Pyr for a week. Cell numbers, determined by counting, were normalized to the glutamine-free control. Statistical comparisons between groups and the glutamine-free control were performed using one-way ANOVA. Data are presented as mean ± SEM from 3 independent replicates. **g**, Relative cell numbers of MDA-MB-231, JIMT-1, MIA PaCa-2, or SUIT-2 after 1-week culture with various supplements: 1 mM α-KG, 200 µM urate or IMP, and 2.5 µM exDNA. Data were normalized to the glutamine-free control and represented as mean ± SEM, n = 3, one-way ANOVA with Dunnett’s multiple comparisons test. **h**, MDA-MB-231 and JIMT-1 breast cancer cells were cultured under glutamine-free conditions for ∼1-week, with or without 25 nM cytD. Treatments included supplementation with 1 mM α-KG or Pyr and 2.5 µM exDNA. Cell numbers were quantified using the CyQUANT assay, normalized to glutamine-free controls, and presented as fold change relative to control (mean ± SEM, n = 6). Statistical significance was determined by one-way ANOVA with Tukey’s multiple comparisons test. **i**, Cell numbers were measured using the CyQUANT assay after 12 days culture in glutamine-deprived medium supplemented with 1 mM α-KG and 2.5 µM exDNA, with or without co-treatment using nucleoside transporter inhibitors (NBTI, DPL, or dilazep) at 0.01 µM (+), 0.1 µM (++), or 1 µM (+++).

Consistent with previous findings^40–42^, we observed that cultured breast cancer and PDAC cells are acutely sensitive to both the GLS inhibitor CB-839 and glutamine starvation (Fig. 2c and Extended Data Fig. 2c). Whereas supplementation with alternative anaplerotic carbon sources (α-KG or pyruvate) rescued proliferation under GLS blockade, it was ineffective at rescuing the effects of glutamine starvation (Fig. 2c and Extended Data Fig. 2c). These observations correspond with the key role of GLS in routing the carbon backbone of glutamine to the TCA cycle, and the broader role of glutamine as a critical source of nitrogen as well as carbon. Supplementing the culture medium with 35–45 mg/mL bovine serum albumin (BSA), comparable to human serum albumin levels^43^, also failed to rescue proliferation under glutamine starvation, indicating that cells cannot obtain sufficient glutamine to sustain growth from the macropinocytic consumption and digestion of extracellular protein (Fig. 2d and Extended Data Fig. 2d). Similarly, combining pyruvate or α-KG with other glutamine-derived metabolites including mixed NEAAs (besides glutamine itself), glucosamine, NAD^+^, or ammonium still failed to rescue proliferation (Fig. 2e). In contrast, when we combined pyruvate or α-KG with mixed deoxynucleotide triphosphates (dNTPs), proliferation under glutamine-free conditions was restored (Fig. 2e).

The *de novo* purine and pyrimidine biosynthesis pathways initially yield inosine monophosphate (IMP) and uridine monophosphate (UMP) respectively, with IMP being a precursor for guanine and adenine nucleotides and UMP a precursor for cytosine, thymine, and uracil nucleotides (Extended Data Fig. 2b). Intriguingly, supplementation with the purine IMP alone was sufficient to rescue growth in glutamine-free media containing an anaplerotic substrate, whereas UMP supplementation did not achieve a significant rescue (Fig. 2f and Extended Data Fig. 2e). This potentially reflects the fact that purine nucleotides are approximately an order of magnitude more abundant than pyrimidine nucleotides in mammalian cells^44^, and/or might reflect detrimental nucleotide imbalance induced by UMP supplementation^45^. The purine urate, which in humans is the end-product of purine catabolism and cannot be salvaged to regenerate nucleotides (Extended Data Fig. 2b), served as a negative control and in contrast to IMP was unable to rescue proliferation (Fig. 2f). Together, these results indicate that anaplerotic substrates such as pyruvate and α-KG can overcome cancer cell dependence on exogenous glutamine-derived carbon, and that an exogenous supply of nucleotides, particularly purine nucleotides, overcomes the requirement for exogenous glutamine-derived nitrogen.

Glutamine depletion is a common metabolic feature of the breast cancer and PDAC TME *in vivo*^46–49^. Our above results show that (i) exDNA is abundant within the TME (Fig. 1a-c), (ii) breast cancer and PDAC cells consume exDNA by macropinocytosis and traffic it to lysosomes (Fig. 1g-j), and (iii) exogenous nucleotides can overcome cancer cell dependence on glutamine-derived nitrogen (Fig. 2d-f). We therefore hypothesized that the macropinocytic consumption and lysosomal digestion of exDNA would decrease cancer cell dependence on *de novo* nucleotide biosynthesis and consequently on exogenous glutamine-derived nitrogen. To test this, we supplied breast cancer and PDAC cells with an anaplerotic substrate together with 2.5 µM exDNA and found that this was as effective as 200 µM IMP at rescuing proliferation under glutamine starvation (Fig. 2g and Extended Data Fig. 2f). The comparable rescue efficacy of multiple randomly generated single-stranded and double-stranded exDNA oligonucleotides indicates that specific DNA sequences are not required for this effect (Extended Data Fig. 2g). In control experiments, we observed that exDNA begins to degrade in culture medium (without cells) after 18 h incubation at 37°C, indicating possible nuclease activity in the dialyzed fetal bovine serum (dFBS) (Extended Data Fig. 2h). We therefore performed high-temperature heat inactivation of dFBS, which successfully eliminated the nuclease activity and thus prevented extracellular breakdown of exDNA even after 3 days incubation at 37°C (Extended Data Fig. 2i), and we found that exDNA still rescues cell proliferation (Extended Data Fig. 2j). Genomic DNA extracted from the non-tumorigenic human mammary epithelial cell line MCF10A—characterized by longer fragments than synthetic DNA along with the presence of epigenetic modifications—was similarly able to rescue cancer cell proliferation under glutamine starvation (Extended Data Fig. 2k).

We next tested if inhibitors of macropinocytosis block the ability of exDNA to overcome cellular dependence on glutamine-derived nitrogen. Treatment with subtoxic doses of cytD or EIPA, which did not impact proliferation under glutamine-replete conditions (Extended Data Fig. 1h,j), sharply suppressed growth when cells were dependent on exDNA as a nutrient source (Fig. 2h and Extended Data Fig. 2l-n). In contrast, blockade of the plasma membrane-localized equilibrative nucleoside transporter SLC29A1 (ENT1) using three structurally distinct inhibitors had no impact on the ability of cells to grow using exDNA, supporting that the observed rescue involves consumption of intact exDNA by macropinocytosis as opposed to extracellular digestion followed by nucleoside uptake (Fig. 2i and Extended Data Fig. 2o).

### Consumed exDNA is digested and maintains cellular purine and pyrimidine pools

Since macropinocytosed exDNA is trafficked to lysosomes (Fig. 1) and can rescue cancer cell dependence on glutamine-derived nitrogen (Fig. 2), we hypothesized that its lysosomal digestion provides a supply of nucleotides that bypasses the need for *de novo* synthesis (Fig. 3a). To test this, we supplied either unlabeled or ^15^N-labeled exDNA, together with α-KG as anaplerotic substrate, to MDA-MB-231 cells deprived of glutamine. For comparison, we grew another set of cells under nutrient-replete conditions containing either unlabeled or [amide-^15^N]-glutamine (Fig. 3b). After 96 h, cells were rinsed, metabolism quenched, and metabolites extracted for analysis by liquid chromatography-mass spectrometry (LC-MS). As expected, in cells grown under nutrient-replete conditions, nucleotide/nucleoside/nucleobase pools were robustly labeled by [amide-^15^N]-glutamine and exhibited the predicted labeling pattern (i.e. m+1 uracil and thymine compounds; m+2 cytosine, inosine, and adenine compounds; and m+3 guanine compounds) (Fig. 3a,c). Remarkably, in glutamine-starved cells supplied with ^15^N-labeled exDNA, ∼75-100% of intracellular nucleotide/nucleoside/nucleobase pools were similarly labeled, indicating that cells had consumed and digested the exDNA to obtain these metabolites (Fig. 3c). Total pool sizes of hypoxanthine and uracil were highly elevated in cells supplied with exDNA relative to those grown with glutamine, indicating that exDNA-derived purines and pyrimidines are processed through nucleotide salvage pathways (Fig. 3a,d). In contrast, early intermediates of the *de novo* purine and pyrimidine nucleotide synthesis pathways such as 5’-phosphoribosyl-N-formylglycineamide (FGAR) and carbamoyl aspartate were labeled as expected by [amide-^15^N]-glutamine but were unlabeled and severely depleted in glutamine-starved cells supplied with exDNA (Fig. 3a,e), consistent with exDNA serving as a source of ‘ready-made’ nucleotides that bypasses the need for *de novo* synthesis.

**Fig. 3:**
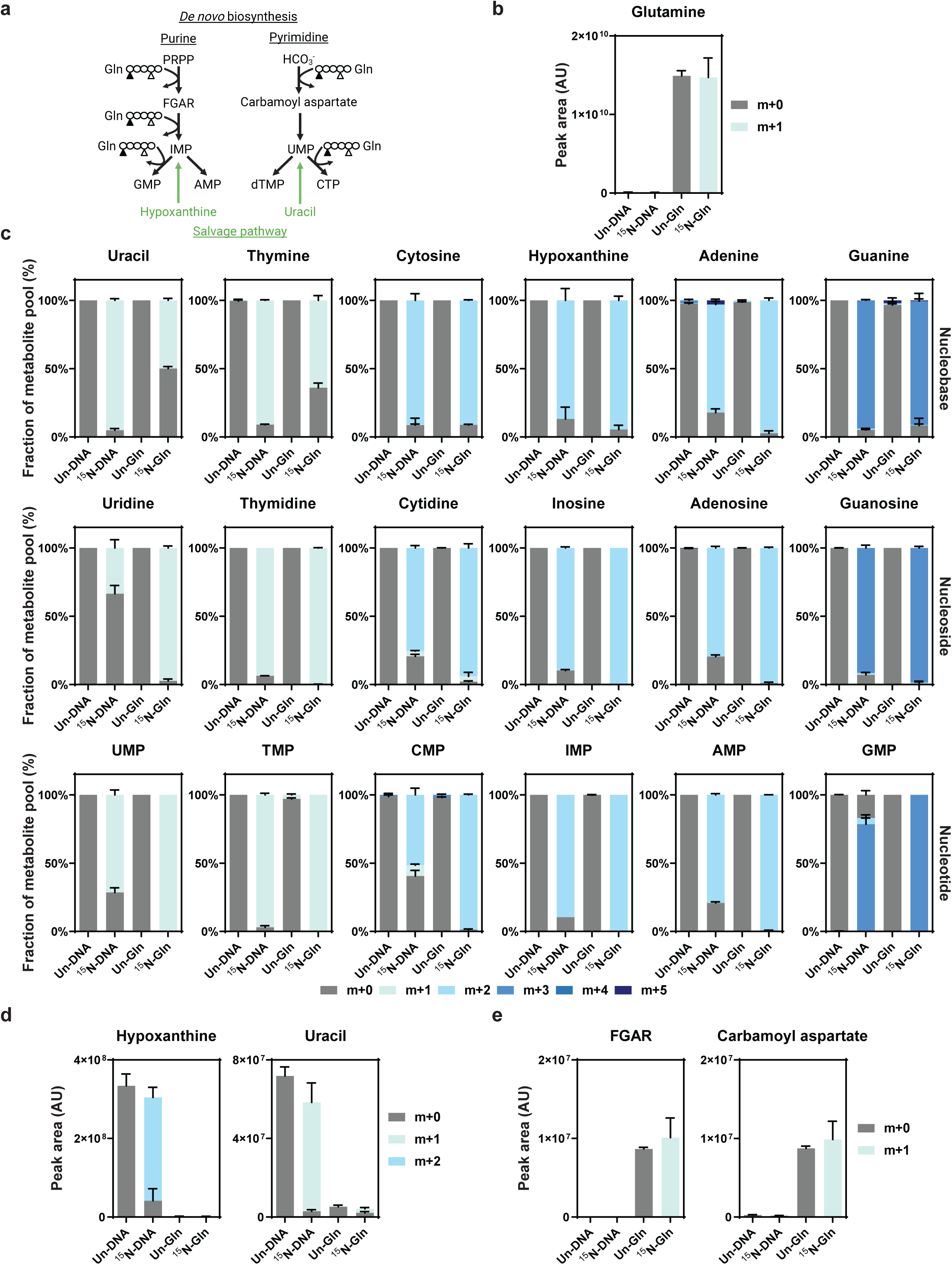
Consumed exDNA is digested and maintains cellular purine and pyrimidine pools. **a**, Schematic illustrating the contribution of nitrogen from [amide-¹⁵N]-glutamine, indicated by black triangles, to key steps in *de novo* purine and pyrimidine synthesis pathways. Salvage pathways are shown in green. **b**, Bar graph showing glutamine abundance and labeling across the samples. **c**, Bar chart showing the proportional isotopologue abundance of nucleobases, nucleosides, and nucleotides following culture with either 1 mM α-ketoglutarate (α-KG) and 23 µg/mL unlabeled or ^15^N-labeled DNA under glutamine-free conditions, or with 2 mM unlabeled or [amide-^15^N]-glutamine. Charts represent the percentage contribution of each isotopologue to the mean overall abundance from 3 biological replicates within each treatment group. **d-e**, Bar graphs showing peak areas from stable-isotope tracing using either ^15^N-labeled DNA or [amide-^15^N]-glutamine, measuring incorporation into nucleotide salvage precursors (d) and early intermediates of *de novo* pyrimidine and purine synthesis (e). Data represent biological triplicates and are shown as mean ± SEM. Values are corrected for natural isotope abundance.

### Use of exDNA as a nutrient source requires lysosomal nucleoside transporter SLC29A3

We next addressed the mechanism by which exDNA is processed to replenish cellular nucleotide/nucleoside/nucleobase pools, following its consumption by macropinocytosis and trafficking to lysosomes. To perturb overall lysosomal function, we first determined cancer cell sensitivity to the lysosomotropic drug chloroquine and then selected a concentration that is subtoxic under nutrient-rich control conditions (Extended Data Fig. 3a). While breast cancer and PDAC cell proliferation in glutamine-replete culture medium was unaffected by this low dose of chloroquine, cells lost the ability to use exDNA to sustain growth during glutamine starvation (Fig. 4a and Extended Data Fig. 3b). Multiple lysosomal nucleases, including DNASE2, DNASE2B, PLD3, PLD4, and ADA2, act on single-stranded and/or double-stranded DNA to yield free deoxynucleotides, which are then converted into deoxynucleosides by a variety of lysosomal 5’-nucleotidases^50–55^. In contrast to the redundancy among lysosomal nuclease and nucleotidase enzymes, the export of (deoxy)nucleosides from the lysosomal lumen into the cytosol is thought to be mediated by a single equilibrative nucleoside transporter, SLC29A3^56^. We therefore tested whether depletion of SLC29A3 impacts the ability of cancer cells to use exDNA as a nutrient source. As with low-dose chloroquine treatment, RNAi-mediated knockdown of *SLC29A3* did not significantly affect the proliferation of breast cancer or PDAC cells under glutamine-replete control conditions (Fig. 4b), but strongly suppressed growth when cells were dependent on exDNA to overcome glutamine starvation (Fig. 4c and Extended Data Fig. 3c). Analysis of gene expression data from The Cancer Genome Atlas (TCGA) and the Genotype-Tissue Expression Project (GTEx) revealed that *SLC29A3* expression is upregulated in breast cancer and PDAC tissue relative to surrounding healthy tissue, a trend that is broadly conserved across multiple cancer types and correlates with higher pathological stages (Fig. 4d and Extended Data Fig. 3d,e)^57,58^.

**Fig. 4:**
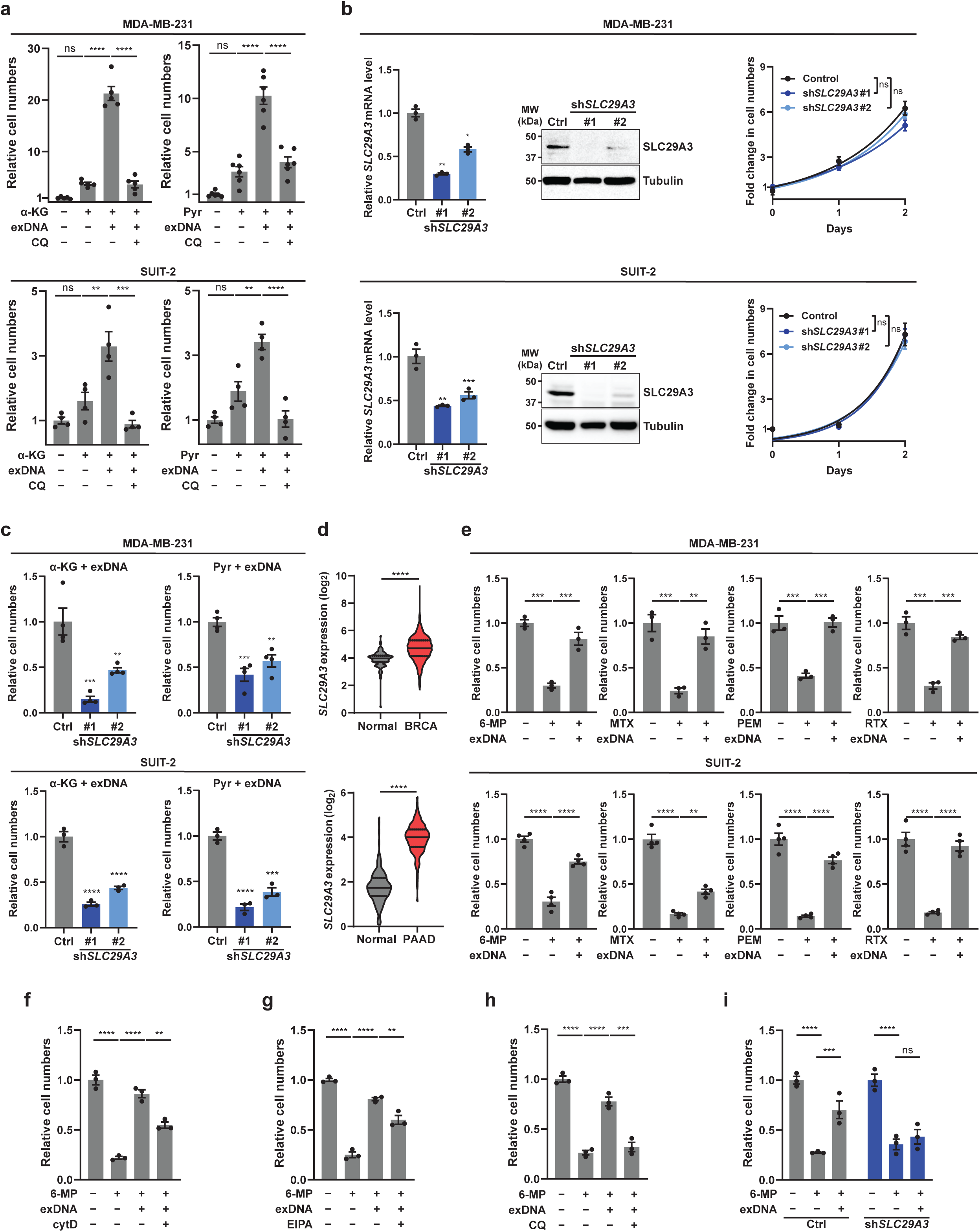
Macropinocytosis of exDNA mediates resistance to nucleotide synthesis inhibitors and requires SLC29A3. **a**, MDA-MB-231 and SUIT-2 cells were deprived of glutamine for 7 days with or without 12.5 µM chloroquine (CQ) for MDA-MB-231 or 25 µM CQ for SUIT-2. Cells were supplemented with either 1 mM α-ketoglutarate (α-KG) or pyruvate (Pyr), along with 2.5 µM exDNA. Cell numbers were quantified using CyQUANT, normalized to glutamine-free controls, and expressed as fold increase above control (mean ± SEM). Statistical significance was assessed by one-way ANOVA followed by Tukey’s multiple comparisons test. **b**, Left: Knockdown efficiency of *SLC29A3* in MDA-MB-231 and SUIT-2 cells assessed by RT-qPCR and western blot. Statistical significance versus control vector was determined by one-way ANOVA with Dunnett’s multiple comparisons test. Right: Proliferation of MDA-MB-231 and SUIT-2 cells following *SLC29A3* knockdown measured by cell counting (mean ± SEM, n = 4). **c**, *SLC29A3* knockdown MDA-MB-231 and SUIT-2 cells were cultured in glutamine-free conditions with supplementation of either 1 mM α-KG and 0.25 µM exDNA or 1 mM Pyr and 0.25 µM exDNA. Cell proliferation was assessed by direct cell counting or crystal violet staining quantification. Values were normalized to the vector control and are presented as mean ± SEM (n = 3). **d**, Gene expression of *SLC29A3* in tumors (TCGA data) and normal tissues (GTEx data) was analyzed using the OncoDB webserver. Violin plots show normal tissue in gray and tumor tissue in red for breast invasive carcinoma (BRCA) (n = 1135 tumor, n = 114 normal) and pancreatic adenocarcinoma (PAAD) (n = 178 tumor, n = 200 normal). **e**, MDA-MB-231 and SUIT-2 cell numbers, normalized to the RPMI control, are presented as mean ± SEM from triplicate experiments after 3 days treatment with 50 µM mercaptopurine (6-MP), 10 µM methotrexate (MTX), 200 µM pemetrexed (PEM), or 15 µM raltitrexed (RTX), with or without 2.5 µM exDNA in RPMI medium. Statistical analysis was performed using one-way ANOVA with Tukey’s multiple comparisons test. **f-h**, MDA-MB-231 cell counts, normalized to the RPMI control, are presented as mean ± SEM from 3 replicates following 3 days treatment with 50 µM 6-MP and 2.5 µM exDNA, with or without 25 nM CytD (f), 12.5 µM EIPA (g), or 12.5 µM CQ (h). Statistical analysis was performed using one-way ANOVA with Tukey’s multiple comparisons test. **i**, *SLC29A3* knockdown suppresses exDNA-mediated 6-MP rescue. MDA-MB-231 cells were treated with 50 µM 6-MP, with or without 2.5 µM exDNA, in RPMI containing heat-inactivated dFBS (70°C, 30 min). Cells were counted on day 3 and normalized to RPMI control.

### Macropinocytosis of exDNA mediates resistance to drugs targeting nucleotide biosynthesis

*De novo* nucleotide biosynthesis is the target of some of the most widely used classes of chemotherapies, including antifolates, antipyrimidines, and antipurines^18,19^. Besides having a direct antiproliferative effect, pharmacological depletion of cellular nucleotide pools sensitizes cancer cells to a spectrum of additional therapies such as genotoxic chemotherapy and radiation^18^. We therefore considered the potential therapeutic implications of our findings. Under nutrient-replete conditions, multiple classes of nucleotide metabolism-targeted drugs, including antimetabolites and non-competitive inhibitors, were effective at suppressing the growth of breast cancer and PDAC cells (Fig. 4e). However, when we supplemented the nutrient-replete culture medium with exDNA, cancer cell proliferation was restored in the presence of each chemotherapy drug, thus rendering these treatments largely ineffective (Fig. 4e). To test if exDNA-mediated chemotherapy resistance requires its macropinocytosis and lysosomal digestion, we blocked this process at different points by applying subtoxic doses of cytochalasin D, EIPA, or chloroquine, or by knocking down *SLC29A3* expression. For this experiment, we focused on the antimetabolite purine synthesis inhibitor mercaptopurine (6-MP), which is a standard-of-care maintenance therapy for some hematological malignancies but has shown only modest efficacy against solid tumors including breast cancer^59–61^. We found that each of these interventions partially or fully resensitizes cells to 6-MP treatment in the presence of exDNA (Fig. 4f-i).

### Targeting SLC29A3 potently sensitizes breast tumors to 6-MP *in vivo*

The results above show that cancer cells can exploit exDNA as a nutrient source to overcome dependence on *de novo* nucleotide biosynthesis. Since the TME *in vivo* is rich in exDNA (Fig. 1)^23,25,27,28^, we hypothesized that blocking a tumor’s ability to process exDNA would sensitize it to chemotherapies targeting nucleotide biosynthesis. To test this, we first determined whether SLC29A3 depletion alone impacts the growth of orthotopic breast tumors in 7-week-old female mice. Whereas *SLC29A3* knockdown had no effect on the proliferation of MDA-MB-231 cells in nutrient-replete culture conditions (Fig. 4b), it modestly but significantly suppressed the growth of MDA-MB-231 tumors *in vivo* (Fig. 5a,b and Extended Data Fig. 4a,b). Real-time quantitative PCR (RT-qPCR) analysis confirmed that *SLC29A3* expression was still robustly depleted in the knockdown tumors at the experiment end point (Extended Data Fig. 4c). To test if SLC29A3 depletion sensitizes tumors to nucleotide biosynthesis-targeted chemotherapy, we again generated control or *SLC29A3*-knockdown MDA-MB-231 orthotopic breast tumors. At ∼1-week post-inoculation, mice bearing control tumors and those bearing *SLC29A3*-knockdown tumors were each divided into 2 groups, with one group then receiving 3 rounds of treatment with 50 mg/kg 6-MP for 5 days on/off and the other group receiving vehicle control. Remarkably, while 6-MP treatment moderately suppressed the growth of the control tumors, it completely blocked the growth of *SLC29A3*-knockdown tumors and even led to significant tumor regression over the treatment course (Fig. 5c,d and Extended Data Fig. 4d).

**Fig. 5:**
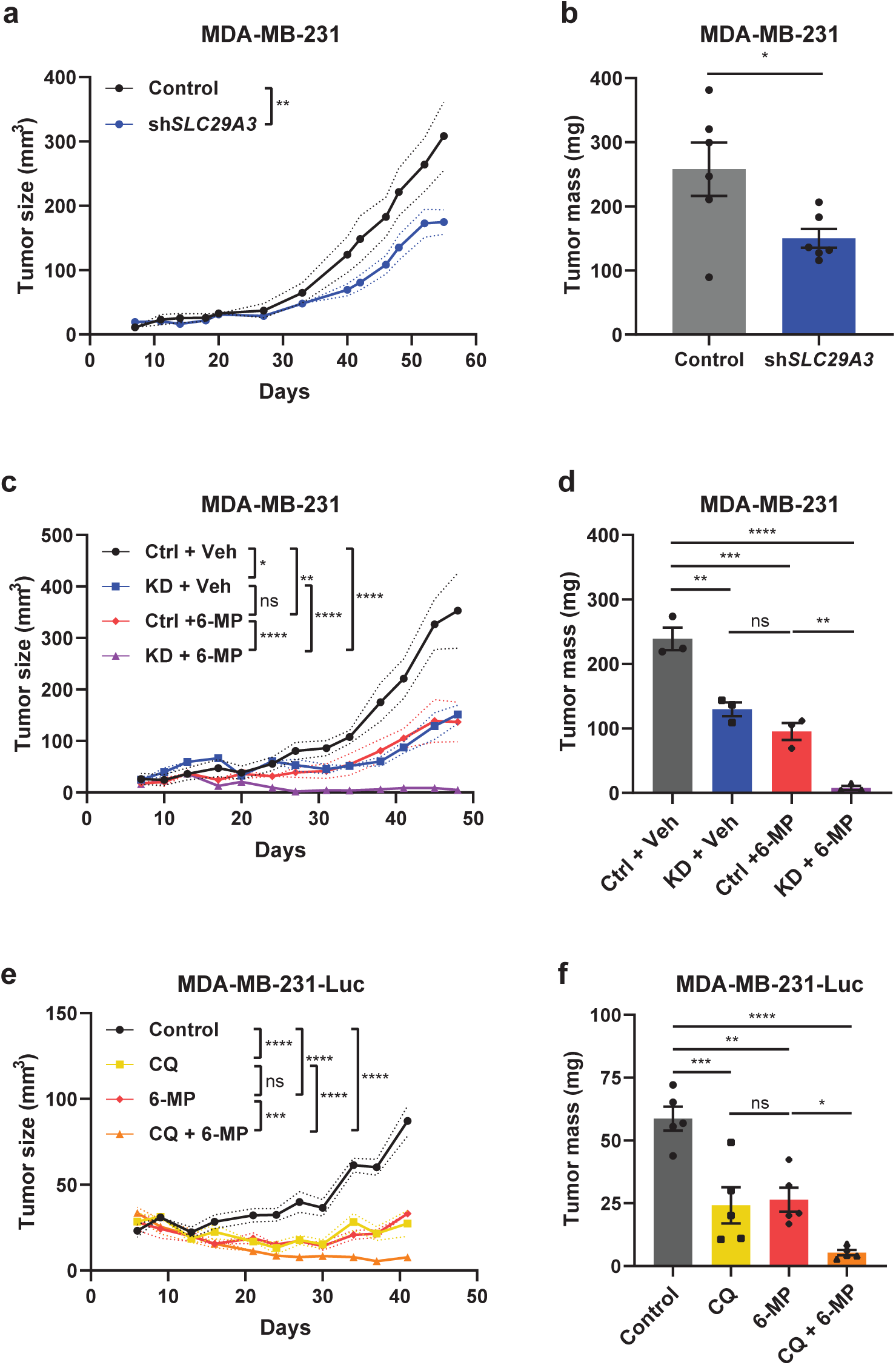
Targeting SLC29A3 sensitizes breast tumors to 6-MP *in vivo* **a**, Volumes of control and *SLC29A3*-knockdown MDA-MB-231 orthotopic xenograft tumors in nude (NU/J) mice. Data are shown as mean ± SEM, n = 6; statistical analysis by two-tailed *t*-test. **b**, Mass of tumors from panel (a), harvested at day 55 post-inoculation. Data are shown as mean ± SEM, n = 6; two-tailed *t*-test. **c**, Volumes of MDA-MB-231 orthotopic xenograft tumors carrying either control vector (Ctrl) or *SLC29A3*-knockdown vector (KD) in nude (NU/J) mice treated with either vehicle (Veh) or 50 mg/kg 6-MP (5-day-on, 5-day-off schedule starting on day 6, for 3 rounds). Data are mean ± SEM, n = 3 per group; two-way ANOVA with Fisher’s least significant difference test. **d**, Mass of tumors from panel (c), harvested at day 49 post inoculation. Data are mean ± SEM, n = 3 per group; one-way ANOVA with Tukey’s multiple comparisons test. **e**, Volumes of MDA-MB-231 orthotopic xenograft tumors in nude (NU/J) mice treated with 50 mg/kg chloroquine (CQ), 50 mg/kg 6-MP (5-day-on, 5-day-off schedule starting on day 6, for 3 rounds), or a combination of both (CQ + 6-MP). Data are mean ± SEM, n = 5 per group; two-way ANOVA with Fisher’s test. **f**, Weights of tumors from panel (g), harvested at day 42 post-inoculation. Data are mean ± SEM, n = 5 per group; one-way ANOVA with Tukey’s test.

Since no selective inhibitor of SLC29A3 has been developed, we considered alternative pharmacological approaches to target the lysosomal processing of exDNA. Chloroquine, which prevents cultured breast cancer and PDAC cells from using exDNA to fuel proliferation during glutamine starvation or 6-MP treatment (Fig. 4a,h), is an FDA-approved drug that has been used to manage malaria, amebiasis, and inflammatory diseases such as lupus^62,63^. We therefore tested whether chloroquine treatment enhances the efficacy of 6-MP against orthotopic breast tumors *in vivo*. Mice were again divided into 4 groups upon detection of palpable tumors and received either vehicle control, 6-MP monotherapy, chloroquine monotherapy, or 6-MP plus chloroquine combination therapy. While each monotherapy moderately suppressed tumor growth, the combination therapy was highly effective and resulted in a steady regression of tumors until the experiment endpoint (Fig. 5e,f and Extended Data Fig. 4e,f).

## Discussion

Autophagy and macropinocytosis are related processes employed by cells to prolong viability under nutrient deprivation stress^64^. Whereas autophagy recycles intracellular substrates, macropinocytosis involves the scavenging of extracellular substrates and thus permits net biomass accumulation^64^. Macropinocytic activity is promoted by some of the most common oncogenic lesions and is frequently activated across a spectrum of malignancies including breast cancer and PDAC^12,13,47,65^. Even in cancer cells with constitutively active macropinocytosis, depletion of certain nutrients including glutamine further stimulates macropinocytic flux^66,67^, and severe glutamine depletion in the tumor core correlates spatially with regions of heightened macropinocytic activity^47–49,66^. In this study, we found that glutamine-deprived breast cancer and PDAC cells can exploit exDNA as a nutrient source by coupling its macropinocytic consumption with a lysosomal digestive pathway that involves the equilibrative nucleoside transporter SLC29A3 (Extended Data Fig. 5). This nutrient acquisition process is sufficient to overcome cancer cell dependence on glutamine-derived nitrogen, which is required for synthesizing purine and pyrimidine nucleotides, and renders chemotherapies targeting *de novo* nucleotide biosynthesis ineffective. Accordingly, genetic or pharmacological disruption of this process abolishes the ability of cancer cells to use exDNA to fuel growth during metabolic stress and potently sensitizes breast tumors to 6-MP.

The macropinocytosis and lysosomal degradation of extracellular protein is known to be an important amino acid supply route in cancer cells in culture and *in vivo*, and can sustain cellular proliferation under conditions of moderate glutamine depletion^16,17,29,30^. Recently, it has emerged that macropinocytic consumption of necrotic cell debris (necrocytosis) provides greater anabolic benefits than protein alone, and can support proliferation under more extreme nutrient stress by supplying a broader range of nutrients including sugars, fatty acids, and nucleotides^65,68^. Our finding that the macropinocytosis of intact exDNA can sustain cellular nitrogen metabolism to fuel proliferation, even during prolonged total glutamine restriction or treatment with chemotherapy inhibitors of *de novo* nucleotide biosynthesis, expands the repertoire of known mechanisms through which macropinocytosis can feed cancer cells within the nutrient-deficient TME. It also suggests that selective pharmacological blockade of critical mediators of this process, such as SLC29A3, could have therapeutic value in enhancing the efficacy of drugs targeting nucleotide biosynthesis, potentially with fewer adverse side effects than broad-action inhibitors of macropinocytosis and lysosomal function.

Our analysis of TCGA and GTEx datasets shows that *SLC29A3* expression is elevated across a spectrum of cancers relative to matched non-cancerous control tissue (Extended Data Fig. 3d). In healthy individuals, *SLC29A3* expression is highly enriched in phagocytes, including macrophages, Paneth cells, Hofbauer cells, Kupffer cells, and trophoblasts^69^. Expression is also elevated in oocytes, which employ a phagocytosis-like pathway to degrade paternal mitochondria after fertilization^70^. The same pathway that we have identified in macropinocytic cancer cells is likely also important for phagocytes to process the vast quantities of DNA that are consumed when dead cells or pathogens are engulfed. Germline loss-of-function mutations in *SLC29A3* are the cause of a variety of familial syndromes that feature abnormal accumulation of phagocytes in multiple organs (histiocytosis) along with associated chronic inflammation^56,71–76^. Recent research has found that this phenomenon involves the sensing of excessive accumulation of nucleosides and oligoribonucleosides by the lysosomal Toll-like receptors 7 and 8 (TLR7/8), which then signal to stimulate monocyte proliferation and proinflammatory cytokine production^74,75,77^.

The presence of extracellular nucleic acids in human blood plasma was first described in the 1940s, and it was subsequently discovered that exDNA is elevated in a variety of pathological states including cancer^23,25,78,79^. In healthy individuals, circulating exDNA levels range from 1–50 ng/mL, whereas in patients with cancer levels can exceed 1000 ng/mL, with the short half-life of ∼15 min indicating sustained and avid exDNA release^25^. Sources of exDNA include cells undergoing apoptosis, necrosis, and NETosis, and the vast majority of exDNA in both healthy and cancer-bearing individuals was recently shown to exist in an accessible form unprotected by biological membranes^28^. Besides entering the circulation, exDNA accumulates within the TME, where its presence is associated with increased metastatic spread and resistance to chemotherapies^26–28,80^.

Although the TME is often deficient in nutrients that are important for cancer cell proliferation, it is rich in macropinocytic substrates that can overcome apparent metabolite addictions, greatly enhancing the metabolic flexibility of tumors. Our study adds exDNA to the suite of TME components that cancer cells can exploit as potential fuel sources, in this case bypassing the requirement for glutamine-derived nitrogen and *de novo* nucleotide biosynthesis. It further suggests that by defining the molecular mechanisms through which a given macropinocytic substrate is processed, promising candidate targets for therapeutic intervention, along with effective new drug combination strategies, can be revealed.

## Methods

### Materials

Chemicals used in this study were as follows: Inosine 5’-monophosphate, IMP (TCI I0036); Uridine 5’-monophosphate, UMP (Sigma U6375); Deoxynucleotide mix, dNTPs (Sigma D7295); Adenosine 5’-triphosphate, ATP (Sigma A6419); Bovine serum albumin, BSA (Sigma A7030); Sodium pyruvate, Pyr (Sigma P2256); Dimethyl alpha-ketoglutarate, α-KG (Sigma 349631); Sodium urate (Sigma U2875); Ammonium chloride (Sigma A9434); N-acetyl-D-glucosamine, GlcNAc (Sigma 1079); D-glucosamine 6-phosphate, Glc-P (Sigma G5509); NAD^+^ (Sigma NAD100-RO); Acetic acid (Sigma A6283); Nucleoside transporter inhibitors: S-(4-Nitrobenzyl)-6-thioinosine, NBTI (Sigma N2255), dipyridamole, DPL (Sigma D9766), and dilazep dihydrochloride (MCE HY-100957). Nucleoside synthesis inhibitors: Methotrexate (Sigma M9929), Pemetrexed (Sigma PHR1596), Raltitrexed (Cayman 26079), and mercaptopurine monohydrate (6-MP) (Sigma 852678). Macropinocytosis inhibitors: 5-(N-ethyl-N-isopropyl) amiloride, EIPA (Cayman 14406) and cytochalasin D, cytD (Cayman 11330). Lysosomotropic agent: chloroquine, CQ (Cayman 14194). Single-stranded and double-stranded DNAs were synthesized by Sigma or Integrated DNA Technologies (IDT). The sequences used were as follows: exDNA #1 and dsDNA: ATCAGTAGAAACTCAGTGTTAGACGATGGA; exDNA #2: TCTTTAGCAGTCAGGTCTATGGAAACTACA. Plasmids for shRNA-mediated *SLC29A3* knockdown were obtained from Sigma (TRCN0000043678, target sequence: GCACAGTTCAAACTCCACCTA and TRCN0000043679, target sequence: CTGTGGCACATACATCATCTT).

### Cell culture

The breast cancer cell lines: MDA-MB-231 and JIMT-1, and the pancreatic cancer cell lines: MIA PaCa-2 and SUIT-2 were cultured in RPMI-1640 medium (Gibco 11875) supplemented with 10% heat-inactivated fetal bovine serum (FBS; Corning 35-010-CV) at 37°C under a 5% CO_2_ atmosphere. For glutamine deprivation, cells were cultured in RPMI-1640 without glutamine (Gibco 21870) and 10% heat-inactivated dialyzed FBS (Hyclone SH30079.03). MCF10A cells were cultured in DMEM/F12 (Gibco 11330) with 5% horse serum (Sigma H1270), 20 ng/mL EGF (Sigma E9644), 0.5 µg/mL hydrocortisone (Sigma H0888), 100 ng/mL cholera toxin (Sigma C8052), and 10 µg/mL Insulin (Sigma I1882). Cell lines were obtained from the American Type Culture Collection (ATCC).

### Cell proliferation and viability assays

Cell proliferation and viability were evaluated through crystal violet staining, cell counting, and the CyQUANT proliferation assay (Thermo C35013). For rescue experiments, cancer cells were plated in complete medium at a density ranging from 20,000 to 50,000 cells per well in a 6-well plate. After 16 h, cells were briefly rinsed and then cultured in glutamine-free media (Gibco 21870) with dialyzed FBS (Hyclone SH30079.03) with indicated concentrations of supplements. The culture media were changed every 2–3 days. Quantification of cell viability was achieved by either de-staining crystal violet with 10% acetic acid and measuring absorbance at OD590 or directly counting the cell numbers (Bio-Rad TC20 or DeNovix). For the CyQUANT proliferation assay, a fluorescence-based DNA content quantification method, cancer cells were plated in 96-well black clear-bottom plates at a density of 2000 cells per well. Cells were then incubated with the indicated drug concentrations, with media changes occurring every other day. Viability was evaluated when the control wells reached confluence in accordance with manufacturer’s instructions. The IC_50_ values were subsequently calculated using GraphPad Prism software.

### Nuclease assays

To validate the specificity of the anti-DNA antibody, 1 µl of 100 µM DNA was treated with or without DNase I (Qiagen 79254) at 37°C for 1 hour. Samples (1.25 µmol) were then mixed with 6x dye and analyzed by tris-borate-EDTA (TBE) 20% polyacrylamide gel electrophoresis. Following electrophoresis, DNA was transferred to a PVDF membrane in 0.5x TBE buffer, blocked with 1% BSA in PBS, and incubated overnight with the anti-DNA antibody (1:200; Novus NBP3-07302). After washing three times with PBS, the membrane was incubated for 1 hour at room temperature with HRP-conjugated anti-rabbit secondary antibody (1:1000; CST 7074). The results were visualized using the Azure Biosystems 300 chemiluminescent western blot imager. To detect potential nuclease activity in FBS 100 ng DNA was incubated in either nuclease-free water, media (Gibco 21870), or media with 10% dialyzed FBS (dFBS) for 1, 2, 4, 6, 18 or 24 hours at 37°C. Separate control tubes contained DNA with DNase I (Qiagen 79254). DNA integrity was analyzed by agarose gel electrophoresis: 4.5% TAE-based agarose gel were stained with ethidium bromide (EtBr) and run at 80 V for 45 minutes, with 100 bp marker (NEB 3231) used for size reference. Gels were visualized using the Azure Biosystems 300 imager under UV302 with auto exposure (∼8 seconds).

### Animal experiments

All animal procedures were approved by the Institutional Animal Care and Use Committee (IACUC) at Cold Spring Harbor Laboratory (CSHL) and conducted in accordance with the NIH Guide for the Care and Use of Laboratory Animals. Six-week-old female nude mice (NU/J, Jackson Laboratory #002019) were acclimated for one week prior to experimentation. Mice were housed in a pathogen-free facility under a 12-hour light/dark cycle, at 20–26°C and 30–50% humidity. Animals received a standard chow diet (PicoLab #5053) and had ad libitum access to food and water. Health status was monitored daily throughout the study. MDA-MB-231 cells carrying either the pLKO.1 control vector or sh*SLC29A3* knockdown vector were orthotopically injected into the fourth mammary fat pads of 7-week-old female nude mice (NU/J, Jackson Laboratory #002019) at a density of 100,000 cells in 0.1 mL PBS with 50% Geltrex (Gibco A1413202). Tumor growth was monitored twice weekly using caliper measurements, and tumor volume was calculated as 0.5 × length × width² with a Vernier caliper. At the study endpoint, tumors were harvested and weighed. In the second experiment, MDA-MB-231 cells expressing either the pLKO.1 control vector or sh*SLC29A3* knockdown vector were orthotopically implanted into the mammary fat pads of nude mice and treated with 6-MP at 50 mg/kg on a 5-day-on, 5-day-off schedule starting on day 5, for three treatment rounds. Tumor growth was monitored by caliper. In the third experiment, 100,000 luciferase-expressing MDA-MB-231 (MDA-MB-231-Luc) cells were orthotopically implanted into the fourth mammary fat pads of 7-week-old female nude mice. Chloroquine (CQ) was administered in drinking water at 0.25 mg/mL, providing an estimated daily dose of 50 mg/kg, starting on day 6. 6-MP was given intraperitoneally at 50 mg/kg in a 5-day-on, 5-day-off schedule starting on day 6, also for three rounds. Tumor growth was monitored using caliper and bioluminescence imaging.

### RNA and DNA extraction and quantitative real-time PCR (RT-qPCR)

Genomic DNA was extracted using QIAamp DNA Mini Kit (Qiagen 51304). Total RNA was isolated from tumor tissue or cultured cancer cells using the RNeasy Mini Kit (Qiagen 74106). Tumor tissues were homogenized with 5 mm stainless beads at 25 Hz for 2 minutes (2 cycles) using a TissueLyser II (Qiagen). The RNA was then treated with DNase I (Qiagen 79254) and cDNAs were synthesized from 1 μg of RNA using SuperScript^TM^ II reverse transcriptase (Thermo Fisher Scientific 18064) with random hexamers following the manufacturer’s protocol. The expression levels of *SLC29A3* and 18S rRNA were quantified using Power SYBR Green PCR Mix (Applied Biosystems) in the QuantStudio 6 RT-PCR system. Primer sequences were as follows: for *SLC29A3*, forward 5’-ATGACCGGCTCCTTTCCTATG-3’ and reverse 5’-GCTGTTCCTCACATCACTGGA-3’; for *18S*, forward 5’-CTGGATACCGCAGCTAGGAA-3’ and reverse 5’-CCCTCTTAATCATGGCCTCA-3’.

### Western blot analysis

Cultured cells were lysed in 1x RIPA buffer (Millipore 20-188) supplemented with proteinase inhibitors (1 µg/mL aprotinin and 1 µg/mL leupeptin) and phosphatase inhibitors (25 mM sodium fluoride and 12.5 mM beta-glycerophosphate) for 15 min on ice. After centrifuge at 10,000 x g for 10 minutes to remove insoluble debris, protein concentrations were determined using a BCA assay. The lysates were then prepared by adding Laemmli buffer (62.5 mM Tris-HCl pH 6.8, 1.5% SDS, 8.3 % glycerol, 0.005% bromophenol blue, and 1.5% freshly added beta-mercaptoethanol). A total of 20 µg protein was resolved on a 4-20% Tris-glycine mini gel and transferred to a PVDF membrane using a semi-dry blot module. The membrane was blocked with 5% non-fat milk in TBST and incubated overnight on a shaker at 4°C with either anti-SLC29A3 antibody (1:1000, Thermo PA5-38039) or anti-tubulin antibody (1:2000, Proteintech 66031-1-1g). After three washes with TBST, the membrane was incubated with anti-rabbit or anti-mouse secondary antibodies at 1:2000 dilution. Following three additional washes with TBST, the signals were developed using Amersham ECL reagents or Pierce™ ECL substrate and imaged using an AZURE 300 chemiluminescent western blot imager.

### Immunofluorescence staining and data analysis

MDA-MB-231 cells were injected into the mammary fat pads of nude mice (NU/J, Jackson Laboratory #002019). Resulting breast tumors were harvested, cryopreserved in Tissue-Tek O.C.T. compound (Sakura 4583), cryosectioned (15 µm) by a Leica CM3050s cryostat, and mounted on Superfrost slides. The tumor slides from the genetically engineered PDAC mouse model (Kras^LSL-G12D^; p53^LoxP^; Pdx1-CreER, KPC), were kindly provided by Dr. David Tuveson (Cold Spring Harbor Laboratory). Human breast and pancreatic cancer tissue blocks were purchased from OriGene Technologies (Sample IDs: FR0001CECE and FR00029545). For DNase I treatment, tumor sections were incubated with 20 µg/mL DNase I (Sigma 11284932001) in 1x DNase I buffer (Thermo B43). Following fixation with fresh 4% paraformaldehyde for 10 min, sections were blocked with 5% goat serum for 1 hour and incubated overnight with primary antibodies targeting DNA (1:200; Novus NBP3-07302) or citrullinated histone H3 (1:1000; Cell Signaling 97272). After washing, sections were incubated with a fluorophore-conjugated secondary antibody (1:500; Thermo A-11008). Nuclear staining was performed with DAPI (1:1000, 1 µg/mL; Thermo 62248) or Hoechst 33342 (1:10,000, 1 µg/mL; Thermo 62249), while cell/tissue boundary staining was performed using CellBrite Fix555 (1:1000; Biotium 30088), Phalloidin labeled PromoFluor 568 (1:500; VWR 10182-884) or Calcein AM (1:500; Thermo C1430). Finally, slides were mounted and imaged using a Zeiss LSM 780 confocal laser scanning microscope equipped with a 63× oil immersion objective, Quasar PMT, GaAsP detectors, and Zen Black 2012 software. In the subsequent experiments, MDA-MB-231, MIA PaCa2, or SUIT-2 cells, seeded on ibiTreat chamber slides (ibidi 80826), were treated overnight with either 2.5 µM Alexa488-exDNA or Cy5-exDNA, with or without co-incubation of DQ-BSA (0.1 mg/mL, Thermo D12050). Cells were then stained for 30 minutes with a combination of Hoechst 33342 (1 µg/mL) for nuclei and, optionally, LysoTracker Red (100 nM) (Thermo L7528) for lysosomes. After PBS washes, cells were maintained in fresh culture media and imaged using the Zeiss LSM 780 confocal microscope.

### Lattice light-sheet microscopy (LLSM) and data analysis

MDA-MB-231 cells were plated onto poly-D-lysine (Gibco A3890401) coated 35 mm MatTek dishes (MatTek P35G-1.5-14-C), incubated at 37°C under a 5% CO_2_ atmosphere, and stained with 100 nM LysoTracker Red (Thermo L7528) for 1 hour prior to imaging. For imaging, we used Zeiss LLSM 7 with 10×/N.A. 0.4 illumination objective lens and 48×/N.A. 1.0 detection objective lens. The illumination light-sheet was 30 μm × 1 μm with no side lobes entering the sample at an angle of about 30° relative to the coverslip. We acquired images in two channels: green channel (DQ-BSA) with excitation at 488 nm and emission at 500–548 nm; red channel (LysoTracker Red) with excitation at 561 nm and emission at 659–720 nm with dual Hamamatsu ORCA-Fusion sCMOS camera configuration. For both channels, we used 10% laser power and 20 ms exposure. For LLSM data processing, we used the Lattice Lightsheet 7 processing module on ZEN blue 3.8 for deskewing, and cover glass transformation. Isotropic pixel size of 145 nm after cover glass transformation. Time-lapse images were recorded every 15 minutes for 16 hours. Further 3D processing and quantitative intensity analysis was then done using Imaris 10.0.1 (Oxford Instrument). The surface function in Imaris was used to create 3D reconstructed surfaces around green (DQ-BSA) or red (LysoTracker Red) microspheres. The surface threshold was set using the automated k-means clustering-based Imaris algorithm. DQ-BSA and LysoTracker Red microspheres were tracked over time using Imaris BT module with the autoregressive motion algorithm to determine macropinocytosis duration for drug treatment timing.

### LC/MS analysis of polar metabolites

MDA-MB-231 cells were seeded in 6-well plates in complete medium and allowed to attach overnight. The culture medium was then replaced with glutamine-free RPMI 1640 medium (Gibco 21870) supplemented with 10% dialyzed FBS (Hyclone 30079.03HI), supplemented with either 2 mM unlabeled glutamine (Sigma G3126), 2 mM [amide-^15^N]-glutamine (Cambridge Isotope Laboratories, CIL, NLM-557-PK), or 1 mM α-KG with 23 µg/mL of either unlabeled or ^15^N-labeled DNA (extracted from cells cultured for >12 doublings in [amide-^15^N]-glutamine). Media were refreshed daily to maintain supplement supply, and metabolites were extracted after 96 hours. Cells were counted from a parallel plate and media then completely removed before adding extraction solution (50:30:20 methanol:acetonitrile:water (v/v/v). After 10 min incubation on dry ice, cells were scraped and transferred into pre-chilled microcentrifuge tubes. Tubes were incubated on dry ice for 1 hour, centrifuged at 13,000 rpm for 10 minutes at 4°C, and supernatants then transferred to new microcentrifuge tubes and lyophilized using a vacuum concentrator. Pellets were reconstituted in 40 µL of 60:40 acetonitrile:water (v/v) plus 10 µL of methanol, vortexing for 20 seconds. Samples were cleared by centrifugation (5 minutes at 4°C), and 10 µL of each supernatant pooled into a clean LC-MS glass vial to serve as the pooled quality control (QC) sample. The remaining volume of each reconstituted sample was transferred to individual clean LC-MS glass vials for analysis. LC-MS analyses for both the polar profiling and isotope tracing experiments were performed using a Vanquish Horizon UHPLC coupled to a QExactive HF mass spectrometer (Thermo Fisher Scientific) via a heated electrospray ionization (HESI) source. Samples were resolved by hydrophilic interaction liquid chromatography (HILIC) using a XBridge BEH Amide column (2.1 × 100 mm, 2.5 µm, Waters) maintained at 40°C. The mobile phases consisted of (A) 20 mM ammonium-acetate in 90:10 water:acetonitrile (v/v) with 0.5% medronic acid, and (B) 2 mM ammonium-acetate in 90:10 acetonitrile:water (v/v) with 0.5% medronic acid. Mobile phases were delivered as follow: from 0 to 12.5 minutes, 100% to 90% B linear gradient (400 µL/min); from 12.5 to 19 minutes, 90% to 60% B linear gradient (400 µL/min); from 19 to 20 minutes, 60% B (400 µL/min); from 20 to 25 minutes, 100% B (800 µL/min). Samples were injected in a randomized order to minimize batch effects, with an injection volume of 5 µL. Eluates were ionized using a HESI source alternating between +3.5 kV to −2.5 kV potential (capillary temperature = 300°C, sheath gas flow rate = 50 a.u., auxiliary gas flow = 10 a.u., and no sweep gas). Ions were measured over an m/z range of 70-700 using at a resolution of 120,000. The automatic gain control target was set to standard with automatic maximum injection time control. Spectral data were processed using FreeStyle software (Thermo Fisher Scientific) and Skyline v.24.1^81^. For both polar profiling and isotope tracing experiments, quantitative analysis was performed on raw peak areas of standard-validated metabolites meeting the following criteria: coefficient of variation less than 30% in pooled quality control samples, signal-to-noise ratio greater than 3, mass accuracy within 5 ppm, and absence of detectable signal in blank controls. In the isotope tracing experiments, raw peak areas were further corrected for natural isotope abundance and tracer impurity using the IsoCorrector R package^82^. These corrected peak areas served as the quantitative basis for downstream analyses. Metabolomics data are deposited in the MetaboLights repository (MTBLS 13024).

## Statistical Analyses

Visualized data and statistical analyses were produced with GraphPad Prism v8.0. Asterisks in the Figure legends indicate significance levels: *, *p* ≤ 0.05; **, *p* ≤ 0.01; ***, *p* ≤ 0.001; ****, *p* ≤ 0.0001; ns, not significant.

## Acknowledgments

We thank all members of the Lukey laboratory and our colleagues in the Demerec building, Dr. Qing Gao, and Dr. Yue Wu for helpful discussions and insights. We thank the Cold Spring Harbor Laboratory Animal, Animal and Tissue Imaging, Mass Spectrometry, and Microscopy Shared Resources, which are funded in part by a National Institutes of Health Cancer Center Support Grant (5P30CA045508). This work was supported by grants from the Department of Defense Breast Cancer Research Program (BC200599), National Institutes of Health (R01GM149957 and 5P30CA045508), METAvivor, Penny’s Flight Foundation, Simons Foundation, and The Elsa U. Pardee Foundation to M.J.L., and from METAvivor to W.-H. Y.

## Author contributions

W.-H.Y. and M.J.L. conceived the study and wrote the manuscript with input from all authors; W.-H.Y., O.T.S., A.S., T.-L.W. performed the experiments; W.-H.Y., A.S., T.-L.W. performed data analysis; E.M., P.C., J.R.C., provided advice on experimental design and data interpretation; M.J.L. supervised the project.

## Extended Data Figures

**Fig. S1:**
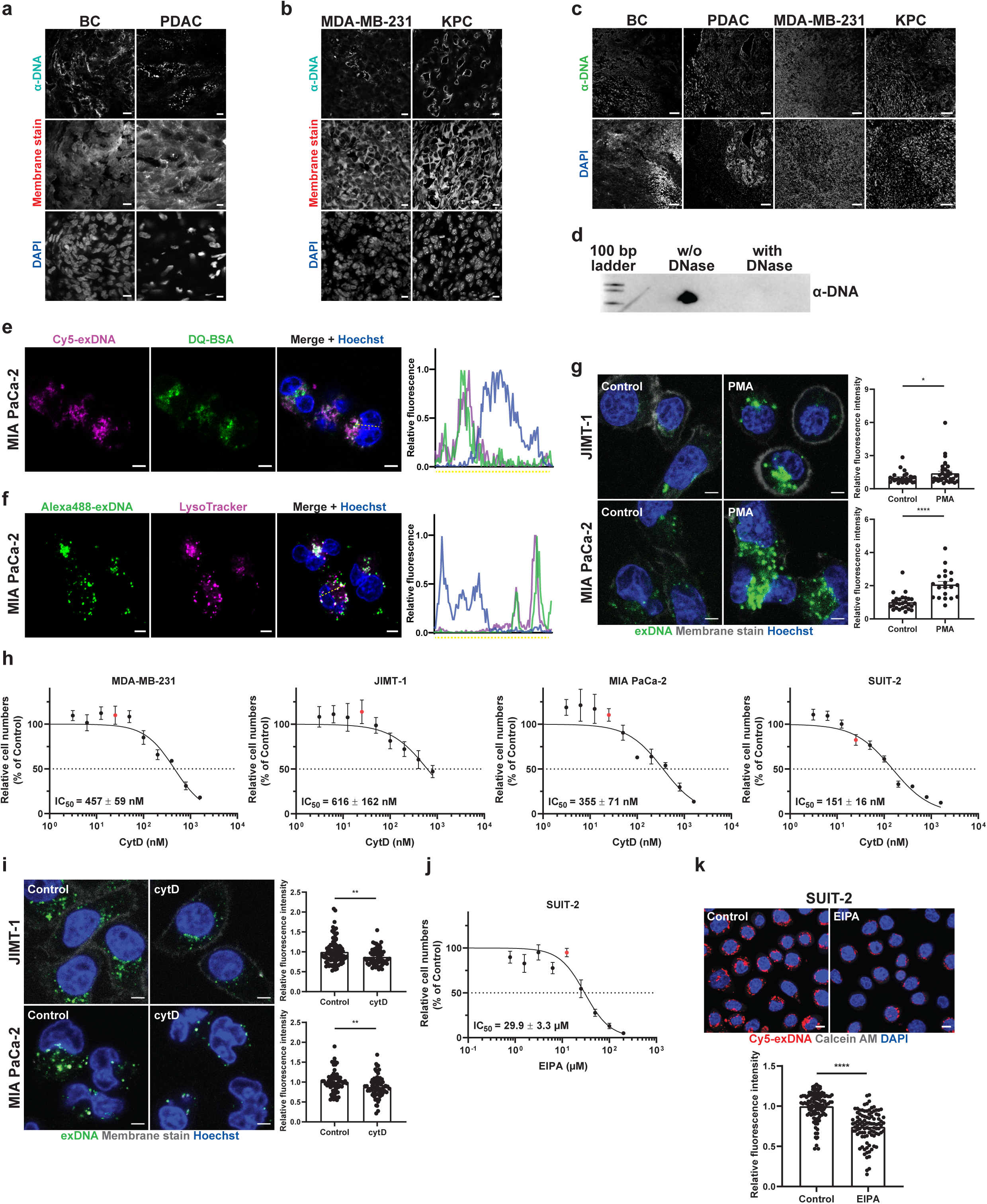
Cancer cells consume exDNA by macropinocytosis and traffic it to lysosomes. **a-b**, Individual channels of confocal microscopy images showing human tumors: Breast cancer (BC) or pancreatic ductal adenocarcinoma (PDAC) in (a), and tumors from mouse models: MDA-MB-231 xenografts and spontaneous PDAC KPC model in (b). Scale bar: 10 µm. **c**, Individual channels of confocal microscopy images of human BC or PDAC, and tumors from mouse models (MDA-MB-231 xenograft and KPC PDAC). Scale bars: 100 µm (BC and KPC) and 200 µm (MDA-MB-231 and PDAC). **d,** Analysis of DNA with or without DNase I treatment using polyacrylamide gel electrophoresis and chemiluminescent western blot. **e-f,** Confocal immunofluorescence of MIA PaCa-2 cells, a PDAC cell line, showing colocalization of exDNA with macropinocytic and lysosomal markers. (e) Cy5-exDNA (magenta) with DQ-BSA (green). (f) Alexa488-exDNA (green) with LysoTracker Red (magenta). Nuclei were stained with Hoechst 33342 (blue). Individual and merged images are shown with corresponding fluorescence quantification plots. Scale bar: 5 µm. **g**, Confocal images of JIMT-1 breast cancer and MIA PaCa-2 PDAC cells treated ± 250 nM PMA and 2.5 µM fluorophore-labeled exDNA overnight, showing exDNA uptake (green). Cell membranes and nuclei were stained with a membrane dye (gray) and Hoechst 33342 (blue), respectively. Scale bars: 5 μm. Vesicle mean intensity was quantified using the Imaris Cell tool. Statistical analysis: two-tailed unpaired Student’s *t* test. **h**, Proliferation of MDA-MB-231, JIMT-1, MIA PaCa-2, and SUIT-2 cells treated with cytochalasin D (cytD), assessed using CyQUANT assay after 3 days treatment in RPMI medium. Data shown as mean ± SEM, n = 4, with IC_50_ values from GraphPad Prism. A concentration of 25 nM cytD is highlighted in red. **i**, Confocal images of JIMT-1 and MIA PaCa-2 cells treated ± 25 nM cytD and Alexa488-labeled exDNA overnight. exDNA uptake (green), membrane (gray), and nuclear (blue) staining as in (g). Scale bars: 5 μm. Quantification and statistical analysis as in (g). **j**, The effect of EIPA on SUIT-2 cell proliferation, assessed using CyQUANT assay after 4 days treatment in RPMI medium. Mean ± SEM, n = 4, with IC_50_ values determined by GraphPad Prism. A concentration of 12.5 µM EIPA is highlighted in red. **k**, Top: Merged confocal immunofluorescence images of SUIT-2 cells with or without 12.5 µM EIPA treatment, showing Cy5-exDNA (red), cell boundary staining by Calcein AM (gray), and Hoechst 33342 (blue) staining of nuclei. Scale bars: 10 μm. Bottom: Quantification of Cy5-exDNA fluorescence intensity was performed by analyzing vesicle mean intensity using the Cell tool in the Imaris analysis platform. Statistical analysis was conducted using two-tailed unpaired Student’s *t* test.

**Fig. S2:**
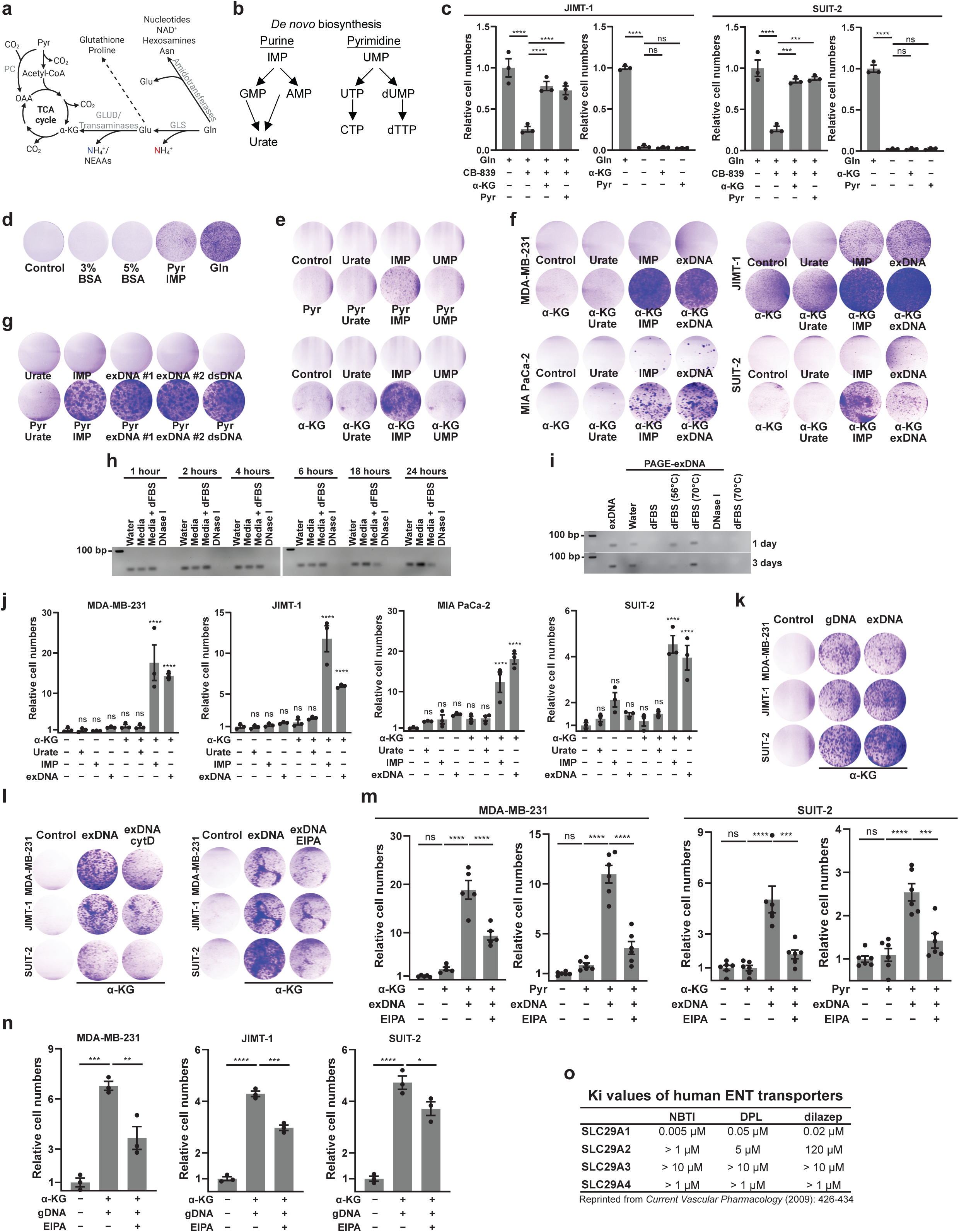
Macropinocytosis of exDNA overcomes cancer cell dependence on glutamine-derived nitrogen. **a**, Summary of glutamine contribution to biomass generation and cellular bioenergetics. Amide (γ) nitrogen is shown in red, amino (α) nitrogen in blue. α-ketoglutarate (α-KG), asparagine (Asn), glutamate (Glu), glutamate dehydrogenase (GLUD), glutaminase (GLS), glutamine (Gln), non-essential amino acids (NEAAs), oxaloacetate (OAA), pyruvate (Pyr), pyruvate carboxylase (PC). **b**, Simplified schematic of *de novo* purine and pyrimidine biosynthesis, including the end-product of purine catabolism in humans, urate. **c**, JIMT-1 breast cancer cells and SUIT-2 PDAC cells were treated for 6 days with CB-839 (10 μM for JIMT-1; 1 μM for SUIT-2) or cultured under glutamine deprivation, with or without 1 mM α-KG or Pyr. Cell numbers were normalized to RPMI control and are shown as mean ± SEM, n = 3. **d**, Representative crystal violet staining of MDA-MB-231 cultured in glutamine-free media supplemented with either 3% or 5% BSA, with 1 mM Pyr plus 200 µM IMP, or with 2 mM Gln for a week. **e**, Crystal violet staining of MDA-MB-231 cells cultured in glutamine-free medium with media changes every 2–3 days. Top, cells were treated with 200 µM urate, IMP, or UMP ± 1 mM pyruvate for 7 days. Bottom, cells were treated with 200 µM urate, IMP, or UMP ± 1 mM α-KG for 10 days. **f**, Representative crystal violet staining of MDA-MB-231, JIMT-1, MIA PaCa-2, and SUIT-2 cells after 1 week of culture in glutamine-free media supplemented with 200 µM urate, 200 µM IMP, or 2.5 µM exDNA, with or without 1 mM α-KG. **g**, Representative crystal violet staining of MDA-MB-231 cells cultured in glutamine-free medium with 2.5 µM exDNA #1, exDNA #2, or double-stranded DNA (dsDNA) ± 1 mM Pyr for 2 weeks. Representative of more than 3 independent experiments. **h**, DNA gel electrophoresis showing nuclease activity against exDNA in dialyzed FBS-supplemented culture media, with DNase I used as a positive control. **i**, DNA gel electrophoresis showing that exposure to 70°C for 30 minutes inactivates the nuclease enzymes present in dialyzed FBS. **j**, Proliferation of breast cancer cell lines MDA-MB-231 and JIMT-1, and PDAC cell lines MIA PaCa-2 and SUIT-2 in a 7-day culture experiment using heat-treated (70°C for 30 minutes) dialyzed FBS. Cell growth was assessed in the presence of 1 mM α-KG, 200 µM urate or IMP, and 2.5 µM exDNA, with glutamine deprivation serving as the control. Cell numbers were normalized to the control and presented as mean ± SEM, n = 3. Statistical significance was determined by one-way ANOVA followed by Dunnett’s multiple comparisons test against the glutamine-deprived control. **k**, Representative crystal violet staining of MDA-MB-231, JIMT-1, and SUIT-2 cells following glutamine deprivation and supplementation with 23 µg/mL genomic DNA extracted from MCF10A cells or 2.5 µM synthetic exDNA, along with 1 mM α-KG, for a week. **l**, Representative crystal violet staining showing that subtoxic doses of macropinocytosis inhibitors cytD (25 nM) (left) or EIPA (5 µM) (right) decrease the rescue effects of 1 mM α-KG and 2.5 µM exDNA from glutamine deprivation in MDA-MB-231, JIMT-1, and SUIT-2 cells. **m**, MDA-MB-231 and SUIT-2 cells were subjected to glutamine starvation for 8 days in the absence or presence of 12.5 µM EIPA, with supplementation of 1 mM α-KG or Pyr and 2.5 µM exDNA. Cell numbers were quantified using CyQUANT, normalized to glutamine-deprived controls, and presented as fold increase above control (mean ± SEM, n = 5 or 6). Statistical significance was determined by one-way ANOVA with Tukey’s multiple comparisons test. **n**, Relative cell density, normalized to the glutamine-deprived control, for MDA-MB-231, JIMT-1, or SUIT-2 cells cultured in glutamine-free medium supplemented with 1 mM α-KG and 23 µg/mL genomic DNA from MCF10A cells, with or without 5 µM EIPA. Data are presented as mean ± SEM, with n = 3. Statistical analysis was performed using one-way ANOVA with Tukey’s multiple comparisons test. **o**, Ki values of human SLC29-family equilibrative nucleoside transporters (ENTs).

**Fig. S3:**
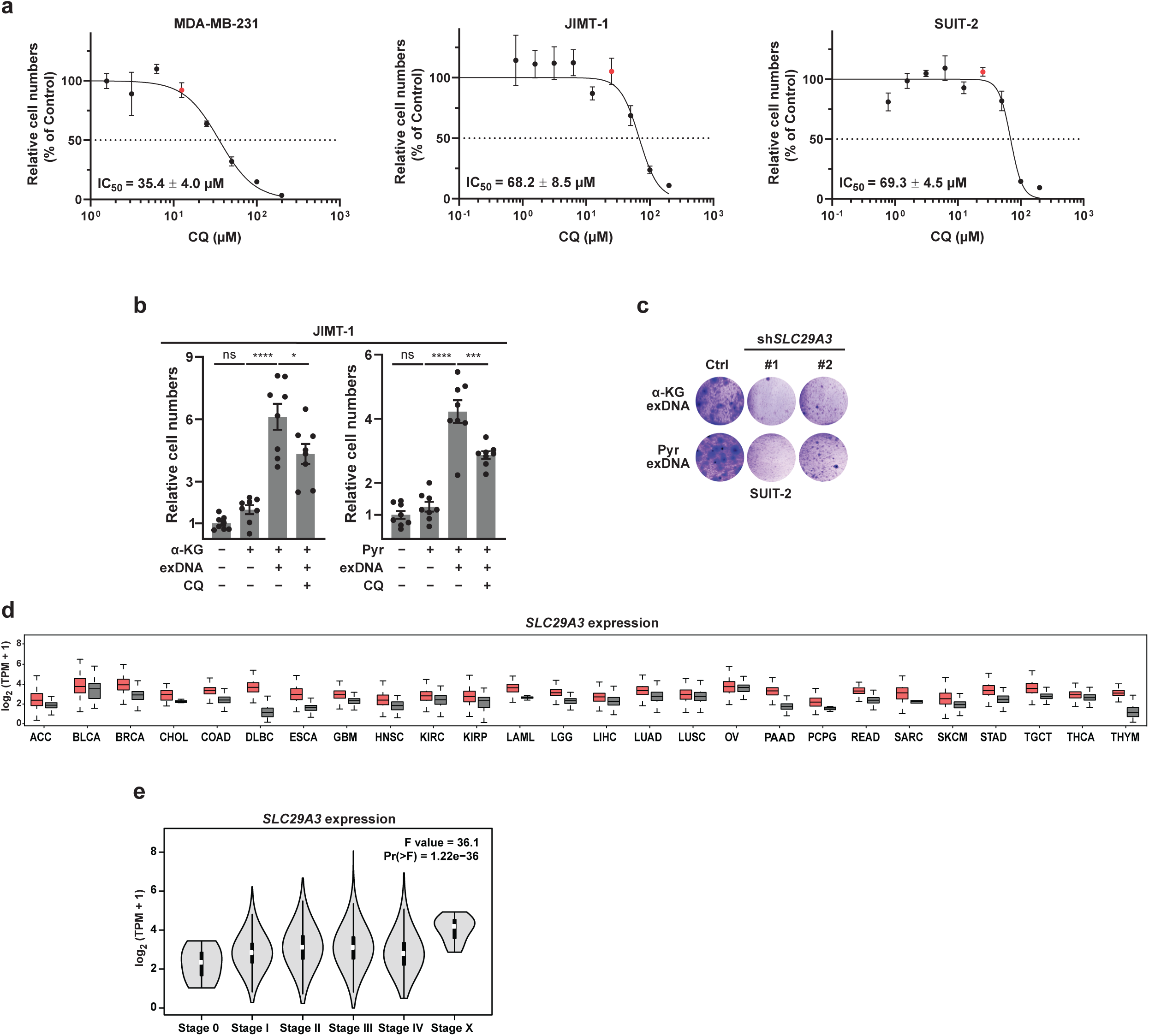
Macropinocytosis of exDNA mediates resistance to nucleotide synthesis inhibitors and requires SLC29A3. **a**, Dose curve for chloroquine (CQ) inhibition of cell proliferation, evaluated using the CyQUANT assay after 4 days treatment. Results are shown as mean ± SEM, n = 4, with IC_50_ values calculated using GraphPad Prism. Highlighted in red are CQ concentrations of 12.5 µM for MDA-MB-231 and 25 µM for JIMT-1 and SUIT-2 cells. **b**, JIMT-1 cell proliferation (5 days) in glutamine-free medium ± 25 µM CQ, with 1 mM α-KG or pyruvate, and 2.5 µM exDNA. Cell numbers were measured by CyQUANT, normalized to glutamine-deprived controls, and expressed as fold change (mean ± SEM). Statistical significance was assessed by one-way ANOVA followed by Tukey’s test. **c**, Effect of *SLC29A3* knockdown on SUIT-2 cell proliferation in glutamine-free medium with 0.25 µM exDNA as nitrogen source and 1mM α-KG/Pyr as anaplerotic substrate. **d**, *SLC29A3* gene expression in human tumors from GEPIA analysis. Gray boxes represent normal tissue, and red boxes indicate tumor tissue in adrenocortical carcinoma (ACC), bladder urothelial carcinoma (BLCA), breast invasive carcinoma (BRCA), cholangiocarcinoma (CHOL), colon adenocarcinoma (COAD), diffuse large B-cell lymphoma (DLBC), esophageal carcinoma (ESCA), glioblastoma (GBM), head and neck squamous cell carcinoma (HNSC), renal clear/papillary cell carcinoma (KIRC/KIRP), acute myeloid leukemia (LAML), lower grade glioma (LGG), hepatocellular carcinoma (LIHC), lung adenocarcinoma/ squamous cell carcinoma (LUAD/LUSC), ovarian serous cystadenocarcinoma (OV), pancreatic adenocarcinoma (PAAD), pheochromocytoma (PCPG), rectum adenocarcinoma (READ), sarcoma (SARC), melanoma (SKCM), stomach adenocarcinoma (STAD), testicular germ cell tumors (TGCT), thyroid carcinoma (THCA), and thymoma (THYM). TPM: transcripts per kilobase million. **e**, *SLC29A3* expression in different pathological stages across all tumor types.

**Fig. S4:**
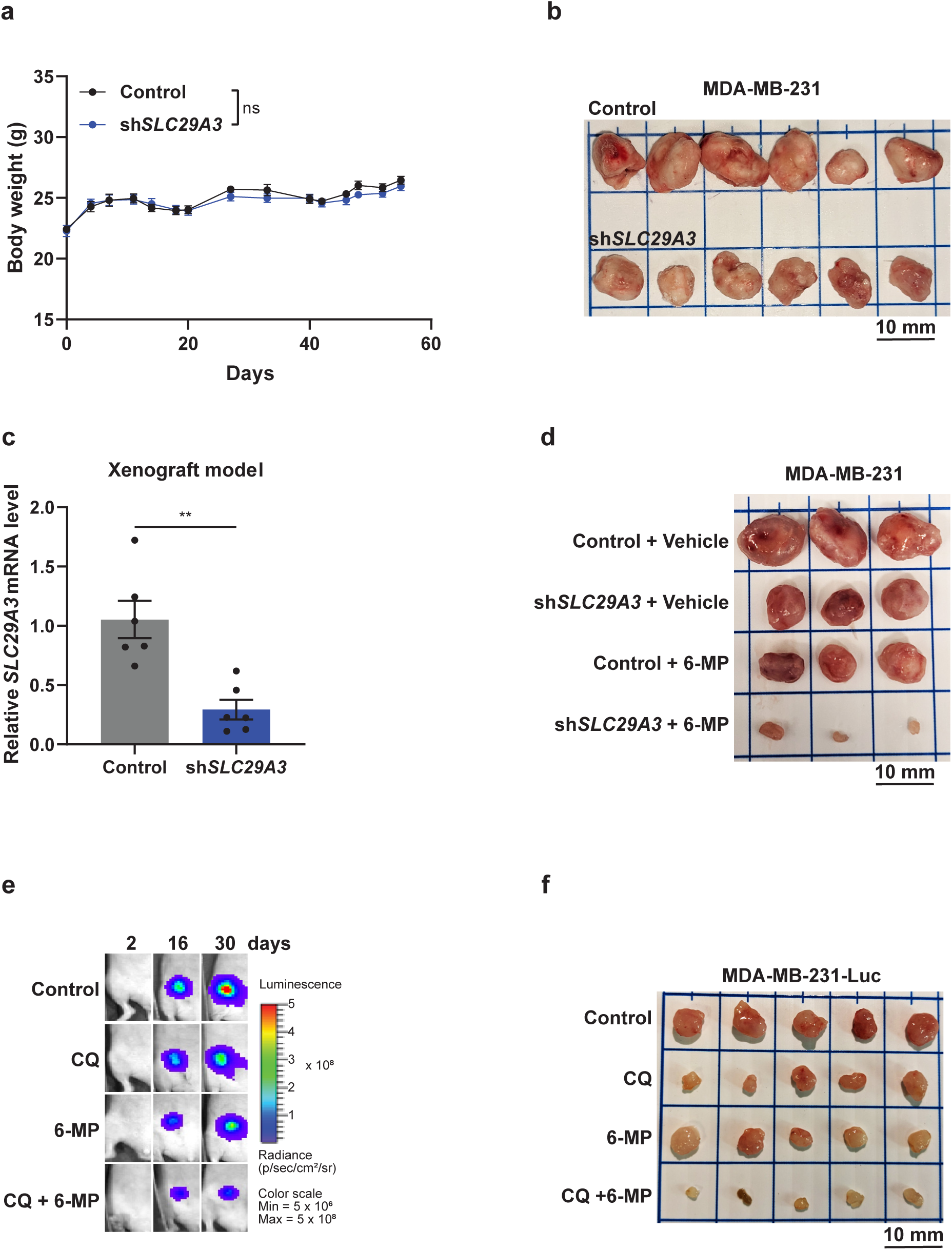
Targeting SLC29A3 sensitizes breast tumors to 6-MP *in vivo* **a**, Body weights of nude (NU/J) mice carrying control or *SLC29A3*-knockdown orthotopic xenograft tumors. Mean ± SEM, n = 6. **b**, Control and SLC29A3-knockdown MDA-MB-231 orthotopic xenograft tumors harvested and imaged 55 days post-inoculation. Scale bar, 10 mm. **c**, RT-qPCR analysis of *SLC29A3* expression in MDA-MB-231 orthotopic xenograft tumors. Data are shown as mean ± SEM with n = 6. **d**, Tumors with SLC29A3-knockdown and/or 6-MP treatment are harvested and imaged at day 49 post-inoculation. Scale bar, 10 mm. **e**, Representative bioluminescence images of MDA-MB-231-luciferase orthotopic xenograft tumors treated with chloroquine (CQ), mercaptopurine (6-MP), or a combination of both, captured on Day 2, Day 16, and Day 30. **f**, Tumors from panel (e) harvested at day 42 post-inoculation. Scale bar, 10 mm.

**Fig. S5:**
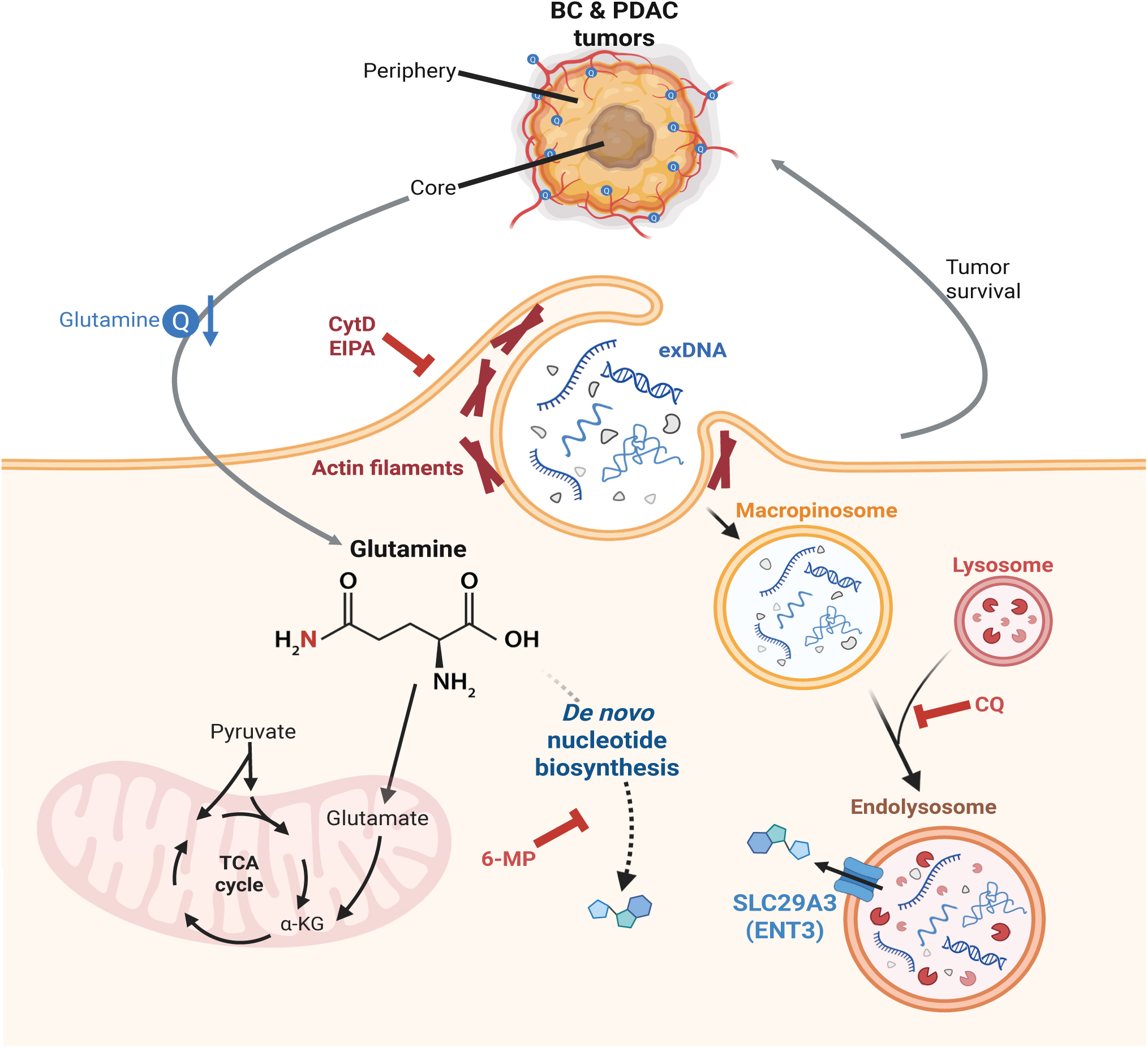
Graphical summary of key findings of this study Created in BioRender. (2025) https://BioRender.com/al2nnjs.

---

**Supplementary Video 1:** Macropinocytosis dynamics in MDA-MB-231 cells Time-lapse sequence of MDA-MB-231 cells labeled with LysoTracker Red (magenta) and incubated with DQ-BSA (green) to visualize intake dynamics during macropinocytosis. Images captured at 15-minutes intervals using ZEISS Lattice Lightsheet 7.

